# Population differentiation and structural variation in the *Manduca sexta* genome across the United States

**DOI:** 10.1101/2020.11.01.364000

**Authors:** Andrew J. Mongue, Akito Y. Kawahara

## Abstract

Many species that are extensively studied in the laboratory are less well characterized in their natural habitat, and laboratory strains represent only a small fraction of the variation in a species’ genome. Here we investigate genomic variation in three natural North American populations of an agricultural pest and a model insect for many scientific disciplines, the tobacco hornworm (*Manduca sexta*). We show that hornworms from Arizona, Kansas, and North Carolina are genetically distinct, with Arizona being particularly differentiated from the other two populations using Illumina whole-genome resequencing. Peaks of differentiation exist across the genome, but here we focus in on the most striking regions. In particular, we identify two likely segregating inversions found in the Arizona population. One inversion on the Z chromosome may enhance adaptive evolution of the sex chromosome. The larger, autosomal inversion contains a pseudogene may be involved in the exploitation of a novel hostplant in Arizona, but functional genetic assays will be required to support this hypothesis. Nevertheless, our results reveal undiscovered natural variation and provide useful genomic data for both pest management and evolutionary genetics of this insect species.

## Introduction

One of the main objectives of evolutionary genetics is to unite the proximate molecular function of genes with the long-term evolutionary forces that govern their change (Nei 1987). The two halves of this pursuit necessitate different approaches, with well-controlled laboratory studies best suited to elucidating the biochemistry of gene function and surveys of natural variation required to understand the ecological and demographic context in which these genes exist. The most fruitful species to study in the lab tend to be those with small body sizes and short generation times. In the field, common species with large population sizes are easiest to sample. Consequently, many of the most popular models for evolutionary genetics come from insects, which fulfill all of the above criteria.

For a brief example, consider the well-studied flies in the genus *Drosophila*. Early pioneers of evolutionary genetics noted that *Drosophila* collected from different locations often had differing arrangements of genes along their chromosomes, in other words, inversion polymorphisms within and between populations (Sturtevant 1917; Dobzhansky and Sturtevant 1938). But without the context provided by population genomic methods, these authors could only catalog the various structural rearrangements they observed; the evolutionary significance of these traits was unknown. With the benefit of modern sequencing technologies, researchers have since shown that segregating inversions like these often exist in environmental clines and contribute to maintaining fitness across a variety of environmental conditions (Anderson *et al*. 2005; Kapun *et al*. 2016; Wellenreuther *et al*. 2017). The initial discovery could only come from controlled laboratory crosses, but the significance could only be established by studying individuals taken from natural populations.

In spite of successes like these, species for which both robust natural and controlled studies exist remain rare. Certainly, the establishment of a new laboratory model species is a massive undertaking and model status can only be achieved through widespread adoption in the research community, which takes years if not decades. Conversely, population genetic sampling is only becoming more tractable with advances in sequencing technology. From this perspective, it is more logical to identify well-studied species that lack population genetic data and bring that natural context to them.

The tobacco hornworm moth (*Manduca sexta*) is a well-established laboratory model (Reinecke, Buckner, & Grugel, 1980), and likely the second most studied moth after the domestic silkmoth, *Bombyx mori* (The International Silkworm Genome Consortium, 2008). The tobacco hornworm is used to study developmental biology of numerous organs (Höglund and Struwe 1970; Banister and White 1987; Nardi 1993), immunology (Kanost *et al*. 2004), reproductive biology (Whittington *et al*. 2015; Mongue and Walters 2017), along with a range of other topics. This attention has resulted in a wealth of genetic information for this species, with many physiological functions characterized by the genetic pathways on which they depend and supported by robust sequencing of RNA (aggregated in Cao and Jiang 2017). In other words, there has been great progress in understanding the molecular genetics of this species.

Yet for all of this research in vitro, the genetics of *M. sexta* in its natural habit remains largely unexplored. Its phylogenetic relationship to other sphinx moths was only recently genetically evaluated (Kawahara *et al*. 2013) and only two years ago were the first whole genome sequences from wild individuals generated (Mongue *et al*. 2019). Very basic questions of natural populations have yet to be addressed. For instance, *M. sexta*, like other moths in the family Sphingidae are particularly strong and mobile fliers (Stevenson *et al*. 1995) and at one time it was claimed that this species is migratory (Ferguson 1991). However, both this initial claim and its later refutation are based solely on limited observations of seasonal occurrence (McNeil 1995) and the ability of pupae to survive subfreezing temperatures (Shapiro 2006). Surprisingly, there have been no genetic studies of population structuring of *M. sexta* to date.

This lack of research into the population genetics of this moth is puzzling on two fronts. First, in terms of basic research, much of the ecological knowledge about this moth, such as acclimation and adaptation to different environments and host plants (Mechaber and Hildebrand 2000; Contreras *et al*. 2013) could benefit from understanding natural genetic variation across its range. Second, as the name tobacco hornworm suggests, *M. sexta* is a pest of economically important crops, including tobacco (*Nicotiana* spp.), tomatoes, peppers, and generally other plants in the nightshade family, Solanaceae (Wagner 2005). Thus, understanding how *M. sexta* populations are structured will inform pest management strategies. In service of both of these goals, we present whole-genome resequencing of this model insect to investigate population structure across an east-west transect of the continental United States. With these resequencing samples, we establish patterns of regional genetic variation and differentiated parts of the genome for further studies of *M. sexta*.

## Methods

### Sample collection and sequencing

For an initial characterization of genetic variation in this broadly distributed species, samples were chosen, based on availability, to maximize the geographic range represented while still sampling local populations with enough individuals to detect large changes in allele frequency. Specifically, samples from three states in the US were examined in this project: North Carolina (n = 12), Kansas (n = 4), and Arizona (n = 8); see Figure 1. Collecting methods varied somewhat between these locales. Samples from North Carolina were already obtained and sequenced as part of Mongue *et al*. (2019). In brief, adults were collected over the course of a week from a mercury vapor light trap in a single locale in July of 2017. Kansas samples were obtained from a mixture of adult moths caught at light traps and larvae taken from tomato fields in different towns in eastern Kansas throughout the summer of 2018. For larval sampling, only one individual was sequenced per field to minimize the risk of sequencing siblings. Arizona samples were obtained from the University of Florida’s collections, where adult moths had been preserved in ethanol. At time of collection, they were collected from a mercury vapor trap at one location over several nights in July of 2012. A summary of sequenced individuals and their methods of collection can be found in Table 1.

**Figure 1.**
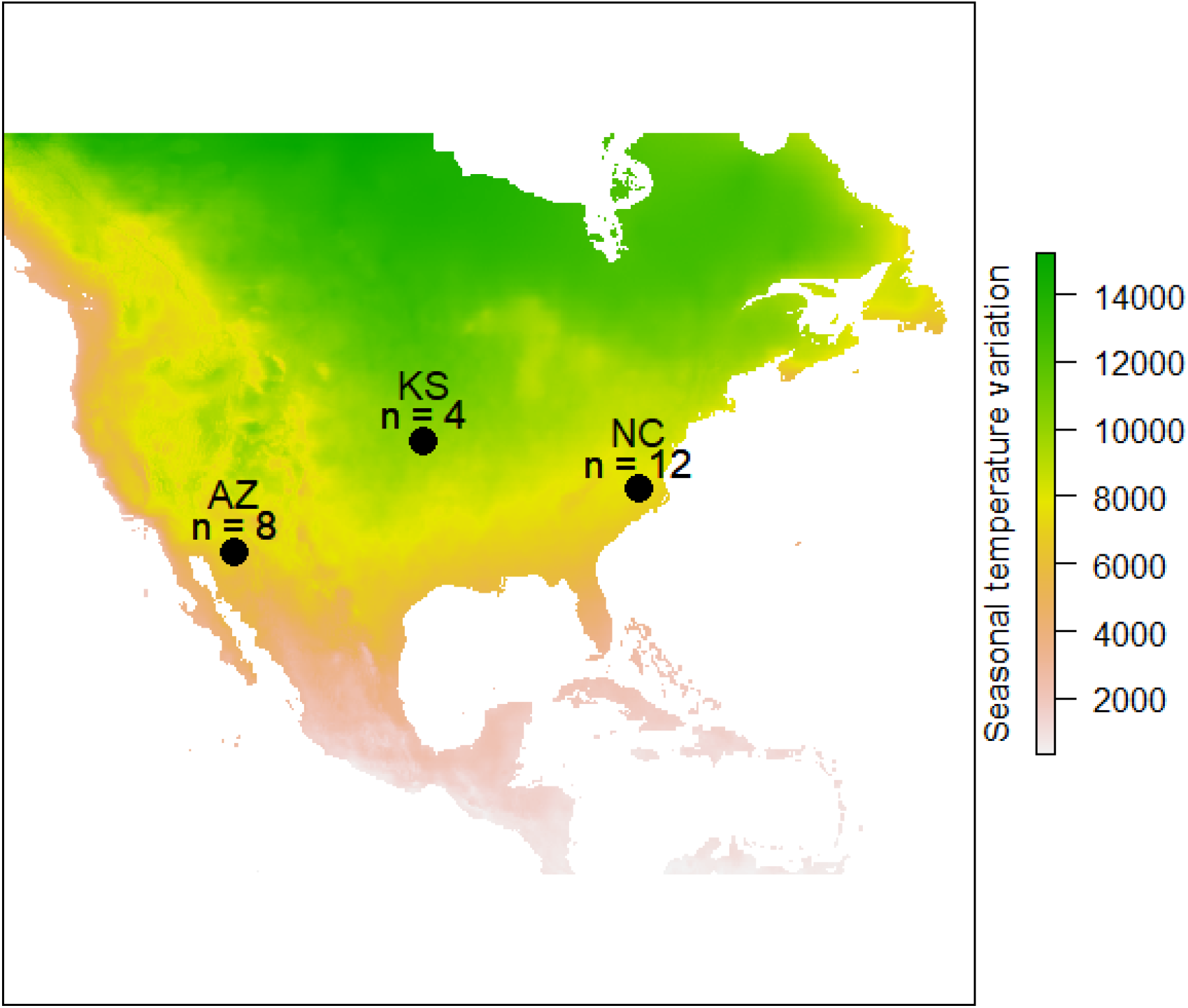
Location of sampled populations of tobacco hornworms with sample sizes. From left to right: Arizona (AZ), Kansas (KS), and North Carolina (NC). Locations are plotted against a backdrop of temperature seasonality (the annual standard deviation in average monthly temperature) to demonstrate that local environment varies across the range of the species.

**Table 1.**
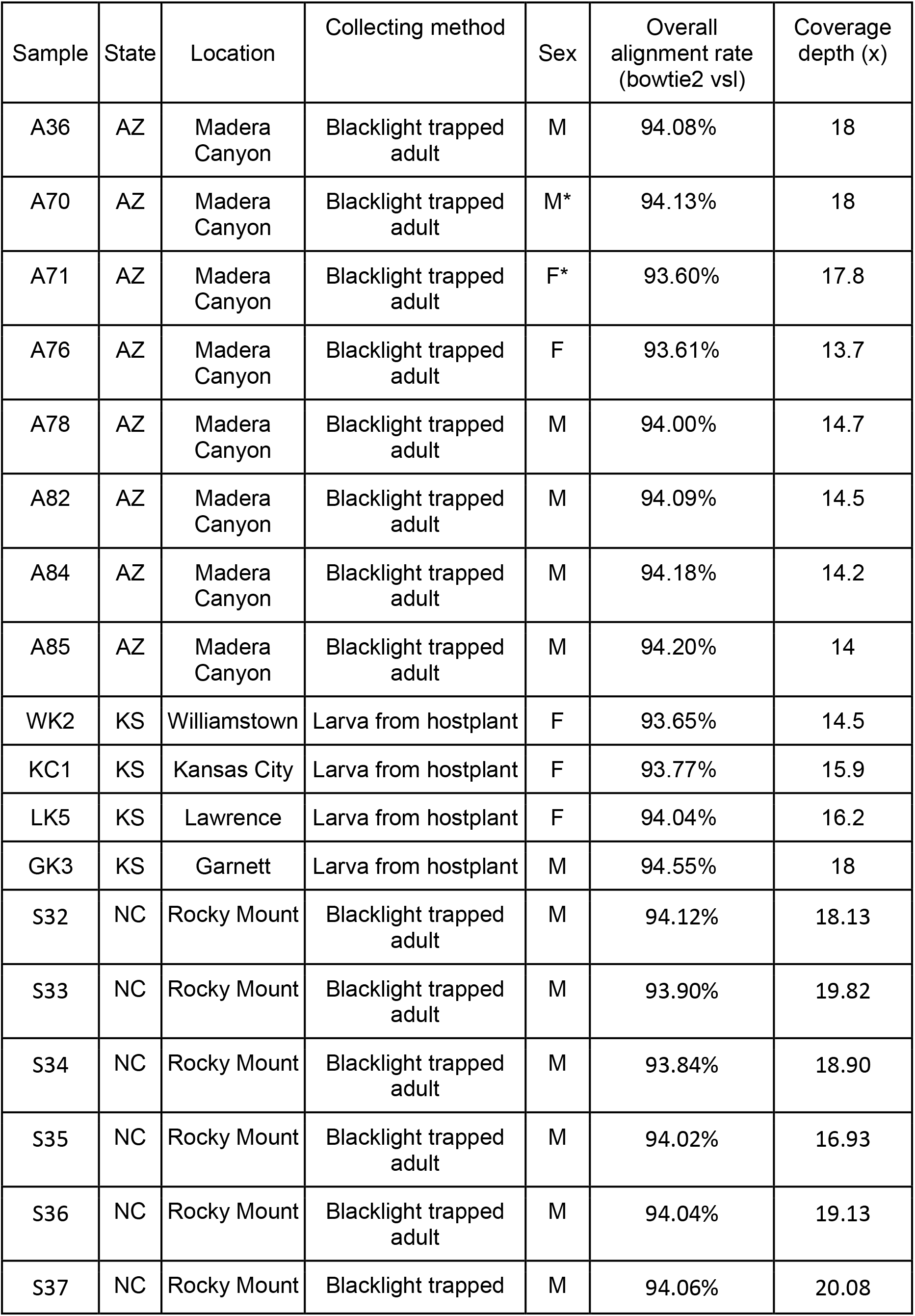
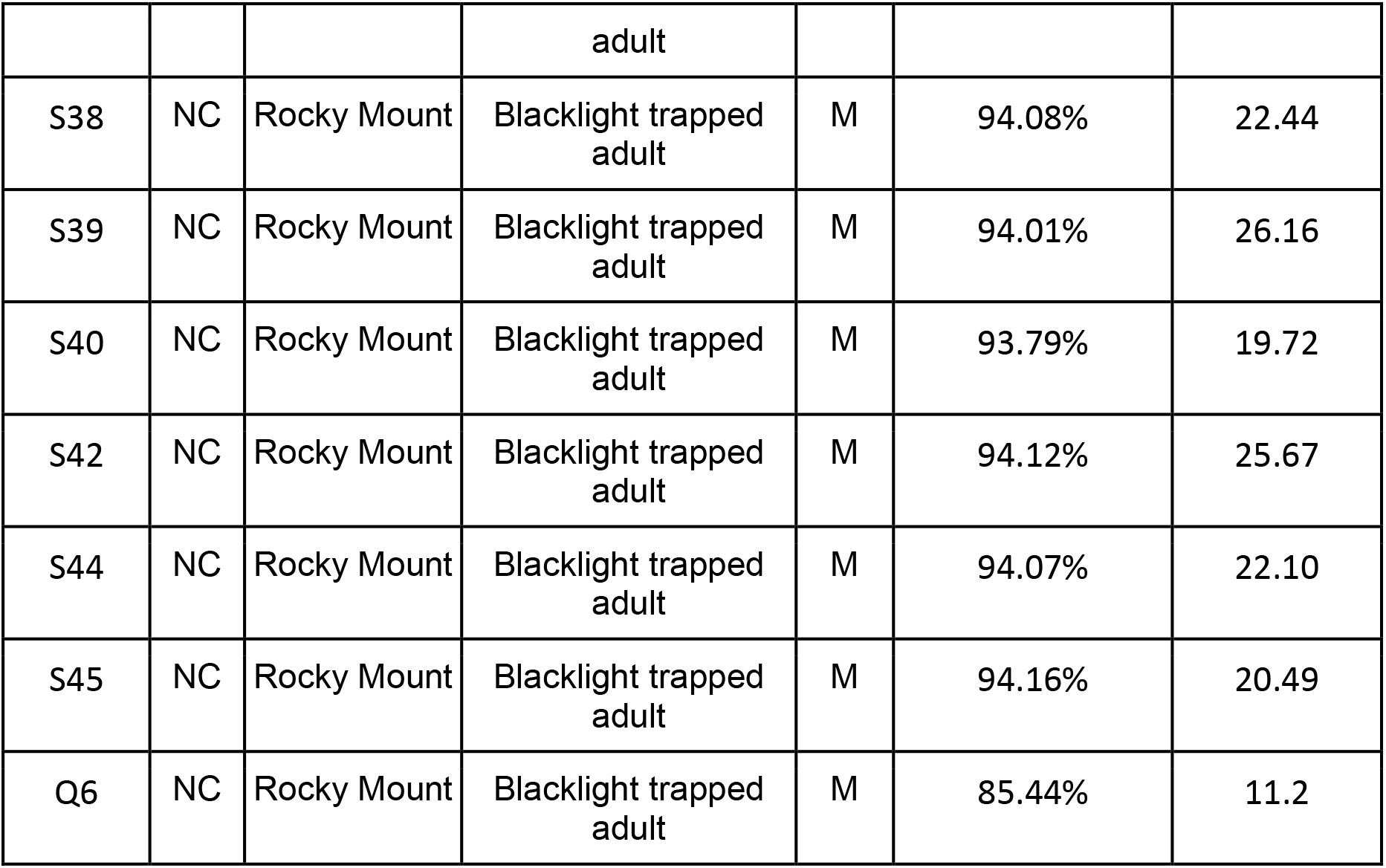
Sample collection and alignment information. Sample names follow sample labeling in SRA. States are abbreviated as AZ (Arizona), KS (Kansas), and NC (North Carolina). Basic genomic information on overall alignment rates to the published *M. sexta* genome (Kanost *et al*. 2016) and depth of sequencing coverage are given as well. All sequencing data are available with the following accessions: SRP144217, PRJNA639154 on NCBI. Individuals with an * next to their sex did not have sex information recorded during sampling; however, we were still able to assign sex based on coverage ratios between the Z chromosome and autosome after sequencing. Q6 is the *M. quinquemaculata* sample used as an outgroup to polarize allele frequencies.

| Sample | State | Location | Collecting method | Sex | Overall alignment rate (bowtie2 vsl) | Coverage depth (x) |
| --- | --- | --- | --- | --- | --- | --- |
| A36 | AZ | Madera Canyon | Blacklight trapped adult | M | 94.08% | 18 |
| A70 | AZ | Madera Canyon | Blacklight trapped adult | M* | 94.13% | 18 |
| A71 | AZ | Madera Canyon | Blacklight trapped adult | F* | 93.60% | 17.8 |
| A76 | AZ | Madera Canyon | Blacklight trapped adult | F | 93.61% | 13.7 |
| A78 | AZ | Madera Canyon | Blacklight trapped adult | M | 94.00% | 14.7 |
| A82 | AZ | Madera Canyon | Blacklight trapped adult | M | 94.09% | 14.5 |
| A84 | AZ | Madera Canyon | Blacklight trapped adult | M | 94.18% | 14.2 |
| A85 | AZ | Madera Canyon | Blacklight trapped adult | M | 94.20% | 14 |
| WK2 | KS | Williamstown | Larva from hostplant | F | 93.65% | 14.5 |
| KC1 | KS | Kansas City | Larva from hostplant | F | 93.77% | 15.9 |
| LK5 | KS | Lawrence | Larva from hostplant | F | 94.04% | 16.2 |
| GK3 | KS | Garnett | Larva from hostplant | M | 94.55% | 18 |
| S32 | NC | Rocky Mount | Blacklight trapped adult | M | 94.12% | 18.13 |
| S33 | NC | Rocky Mount | Blacklight trapped adult | M | 93.90% | 19.82 |
| S34 | NC | Rocky Mount | Blacklight trapped adult | M | 93.84% | 18.90 |
| S35 | NC | Rocky Mount | Blacklight trapped adult | M | 94.02% | 16.93 |
| S36 | NC | Rocky Mount | Blacklight trapped adult | M | 94.04% | 19.13 |
| S37 | NC | Rocky Mount | Blacklight trapped adult | M | 94.06% | 20.08 |
| S38 | NC | Rocky Mount | Blacklight trapped adult | M | 94.08% | 22.44 |
| S39 | NC | Rocky Mount | Blacklight trapped adult | M | 94.01% | 26.16 |
| S40 | NC | Rocky Mount | Blacklight trapped adult | M | 93.79% | 19.72 |
| S42 | NC | Rocky Mount | Blacklight trapped adult | M | 94.12% | 25.67 |
| S44 | NC | Rocky Mount | Blacklight trapped adult | M | 94.07% | 22.10 |
| S45 | NC | Rocky Mount | Blacklight trapped adult | M | 94.16% | 20.49 |
| Q6 | NC | Rocky Mount | Blacklight trapped adult | M | 85.44% | 11.2 |

For each sample, tissue was extracted from adult thoracic flight muscle to avoid eggs and sperm in the abdomen, and larval head capsules to avoid hostplant tissue in the gut. DNA was extracted using an Omniprep kit for genomic DNA (G-Biosciences, St. Louis, MO, USA), following the manufacturer’s protocol. DNA was isolated via phase-separation with chloroform and precipitated with ethanol. DNA pellets were resuspended in buffer and sent for Illumina sequencing with Novogene Sequencing (Sacramento, CA, USA). Quality control and library preparation followed the company’s standard and sequencing generated 150 basepair, paired-end reads sequenced to an average depth of 14-20x coverage across the genome (also summarized in Table 1). As part of the sequencing service, the company trimmed raw reads for both quality and adapter sequence content.

### Starting points for genomic analyses

Recent work on the *Manduca sexta* genome assembly has created both the opportunity and need for some comparative work on the multiple available assemblies. *Manduca sexta* has had an Illumina-only (i.e. fragmented) genome assembly for several years (Kanost *et al*. 2016); much more recently a chromosome-level assembly has been published (Gershman *et al*. 2021). Naturally, the chromosome-level assembly’s more contiguous scaffolds facilitate identification of large-scale variation; however, the previous assembly features high quality gene models informed by dozens of RNA-sequencing experiments and manually updated over the last five years (Kanost *et al*. 2016). To take advantage of the strengths of both assemblies, the same analysis pipeline was carried out on both assemblies to leverage the wealth of available data. Generally speaking, the newer assembly was employed for genome-wide investigation of differentiation and the older assembly was used to validate patterns observed in the former and investigate specific gene(set)s of interest. For both assemblies, the same processed short-read data were aligned to the reference using the very-sensitive-local alignment setting in Bowtie2 version 2.2.9 (Langmead & Salzberg, 2012). Alignments were coordinate-sorted and optical duplicates were removed with Picardtools v. 2.8 (Wysoker *et al*. 2013). Finally, alignments around insertions and deletions were adjusted with GATK v. 3.7 (McKenna et al., 2010). This end result generated a set of curated alignment files (bams) that were the starting point for all downstream analyses.

For some of these analyses, it was necessary to analyze the genotypes of single nucleotide variants. To generate sets of high-confidence variants, the curated alignments were taken through GATK v 3.7’s best practices pipeline for SNP calling (McKenna *et al*. 2010), including hard quality filters (Quality by Depth > 2.0 & Fisher Strand-bias < 60 & Mapping Quality > 40). The end result was a Variant Call Format (vcf) file containing SNPs for each population. This filtering resulted in a total of 16,674,625 SNPs across all of the *M. sexta* samples on the new assembly (and a similar 16,792,392 for the old assembly).

Finally, some inferences of population differentiation relied on differences in allele frequency between locations. For polymorphic sites, estimation of allele frequency depends on inference of which of the alleles (reference or alternate) is ancestral, otherwise allele frequencies are collapsed to the range [0.0, 0.5] and information is lost. To infer ancestral state at a given site, variation in *M. sexta* was compared to that in a congener, *Manduca quinquemaculata*. A *Manduca quinquemaculata* sample was sequenced in Mongue *et al*. (2019). Reads from that individual were aligned to the references with Stampy v1.08 (Lunter and Goodson 2011), using “--substitutionrate=0.1” to allow for increased single nucleotide mismatches expected from divergences. After initial alignment, this sample was treated in the same manner as the within-species samples (curated with Picard and variants called with GATK), producing 9,333,270 high quality single nucleotide variants. These putative divergences were combined with the polymorphism data from *M. sexta* to create a new “ancestral” reference (fasta) for use in polarizing allele frequencies, using the following parsimony logic.

In a majority of cases, polymorphic sites did not overlap with divergent sites as well. In other words, these sites that varied in *M. sexta* but not in *M. quinquemaculata* showed the reference allele in the outgroup; thus, the reference allele was taken to be ancestral and left as was. For each site that varied in both *M. sexta* and *M. quinquemaculata,* allele identities were compared between species. If the alternate allele in *M. quinquemaculata* was shared with the alternate allele in *M. sexta* the alternate allele was inferred to be ancestral, and the reference derived. Likewise, if the outgroup was heterozygous at the site, but shared only one allele with the focal species, that allele was inferred to be ancestral (e.g. a tri-allelic site with reference: T, *M. sexta* alternate allele: C, *M. quinquemaculata* alternate alleles: G, C). Considering both of these scenarios, a total of 1,327,986 sites were updated in the reference. In a minority of cases (e.g. a tri-allelic site with reference: T, *M. sexta* alternate allele: C, *M. quinquemaculata* alternate allele: G), there was no most-parsimonious solution. These sites were updated to “N” at the site in the new ancestral reference to mask them from analysis (n = 275,410 sites). The code to generate the updated reference was written in R and can be found in the git repo listed in Data Accessibility.

### Updating genomic resources

In addition to preparation of resequenced samples and variants, the newer reference genome required ancillary metadata, namely: the assignment of scaffolds to chromosomes to separate chromosome-specific effects (e.g. differing dynamics on the Z sex chromosome). Curiously, the newer *M. sexta* reference contains no chromosomal linkage information for its scaffolds. The older assembly does have linkage information however, from previous ortholog counting analyses comparing *M. sexta* to *B. mori* to identify and separate sex-linked scaffolds; this effort generated a chromosome identity for each scaffold, but did not contain information about the ordering of scaffolds within a chromosome (Mongue and Walters 2017). To anchor the newer assembly, it was aligned to the older one using SatsumaSyntenty in Satsuma v.3.1 (Grabherr *et al*. 2010). This tool generated a list of alignments between the older, anchored assembly and the newer, more contiguous but unanchored assembly. Chromosomes were assigned to the new assembly’s scaffolds by tabulating the number of fragmented (older) scaffolds aligning to each chromosome-length scaffold and taking the highest scoring chromosome. This effort generated unambiguous assignment of the largest scaffolds in the newer assembly (which should be mostly complete chromosomes) as well as putative assignments for the other, unplaced scaffolds. These placements were mostly unambiguous based on syntenic alignments alone, but in the case of two ambiguities, additional information from BAC-FISH probes between *M. sexta* and *B. mori* were used to resolve chromosomal assignment (Yasukochi *et al*. 2009). As this information may be of future use beyond the scope of this research, both this assignment and the alignment summary file used to generate it can be found in the supplementary files.

### Population structure

With linkage established, alignments were filtered to separate out the Z chromosome, as unequal sampling of sexes between populations (e.g. North Carolina’s sample is male-only) could create spurious patterns. The autosomal genome was used to infer population structure while accounting for uncertainty in genotyping using the software NGSAdmix v. 3.2 (Skotte *et al*. 2013) as well as a principle-component based approach with PCAngsd v.0.98 (Meisner and Albrechtsen 2018). For the admixture analysis, the best-supported number of populations (K) was determined by assessment of ten bootstrapped replicates each for possible K values from 2 to 6, evaluated using the logic from CLUMPAK’s web-hosted application (Kopelman *et al*. 2015). After a first round of analyses revealed a large (~8Mb) block of differentiation on chromosome 12 segregating in population, structuring analyses were repeated with this chromosome removed as well.

### Characterization of genomic differences

After establishing basic population relationships, population differences were examined in detail. A baseline expectation of differentiation was set by estimating the genome-wide pairwise F_ST_ (excluding the Z chromosome) analyzing the curated alignments with the population genetic software ANGSD v. 0.916 (Korneliussen *et al*. 2014). In more detail, we input into angsd the list of curated bam files and the parsimony-informed ancestral reference sequence and generated site allele frequency likelihoods (-dosaf 1) separately for each population using the SAMtools method of genotype likelihoods (-gl 1). Next, a two-dimensional site frequency spectrum was generated for each pair of populations for a total of three comparisons using the realSFS command. With the relevant site frequency spectrum as a prior, we used the -fstout command to generate genome-wide F_ST_ on using both assemblies as references. Additionally, we estimated F_ST_ separately for silent (four-fold degenerate) and non-silent (zero-fold degenerate) sites to assess whether differentiation skewed toward silent sites or those more likely to be under selection. Identification of zero-fold and four-fold degenerate sites was completed in previous analyses with custom R scripts (in R v 3.3, R Core Team 2017; Mongue *et al*. 2019) but was only carried out on the Kanost et al. assembly due to the higher quality gene models.

With a baseline established, candidates for further investigation were identified via a preliminary scan for F_ST_ outlier regions in windows of 10 kilobases with a 5 Kb step size to identify regions for further study (using the fst command again with -win 10000 -step 5000). All pairwise comparisons were performed: North Carolina (NC) - Arizona (AZ), Kansas (KS) - Arizona, and NC - KS. For these analyses, published chromosome assignments of scaffolds were employed for the older assembly (Kanost *et al*. 2016; Mongue and Walters 2017; Mongue *et al*. 2020) and assignments generated for this manuscript were used for the new assembly. For the older assembly, these assignments did not contain information on ordering of scaffolds *within* a chromosome. Thus, we focused primarily on the new assembly for identification of differentiated regions, but we performed analogous F_ST_ analyses with alignments to the old assembly to assess differentiation in and around coding regions.

These analyses revealed multiple large F_ST_ peaks, as would be expected of segregating inversions. To further investigate these regions and characterize which population(s) carried the ancestral or derived orientation in the putatively inverted regions, genetic diversity was examined as follows. Single nucleotide polymorphism (SNP) frequencies were examined in each population, under the logic that if loci within an inversion cannot freely recombine with the non-inverted orientation, they should share allele frequencies across longer distances than those outside of the inversion. For these analyses the polarized allele frequencies described above were used. SNP variant files (vcfs) were generated for each population and alternate allele frequencies were extracted and examined. Beyond allele frequency, the allele identity was also examined. The VCFtools utility “extract-FORMAT-info GT” (Danecek *et al*. 2011) was used to examine sample genotypes directly and determine which individuals carried non-reference alleles across genetic features of interest. Finally, to further evaluate the interpretation of these regions as inversions, a pair of individuals ostensibly carrying the inversions was investigated with Delly2 v 0.8.7 (Rausch *et al*. 2012) to identify structural variants and potential inversion breakpoints. Because Delly makes inferences based on read-pair alignment positions, the individuals with the deepest coverage of those carrying the potential inversions were chosen (A36 and A70) to compare to a pair ostensibly lacking the inversion (S39 and S42).

Finally, in the interest of establishing baseline metrics for these populations, we calculated a number of population genetic summary statistics. In particular, we calculated measures of variability: pN (nonsynonymous changes per nonsynonymous site), pS (synonymous changes per synonymous site), pN/pS the scaled ratio of non-synonymous variation, π_0_ (variation at zero-fold degenerate sites), π_4_ (variation at four-fold degenerate sites), Tajima’s D_0_, and Tajima’s D_4_ (estimators of deviation from neutral evolution at zero- and four-fold degenerate sites respectively). These metrics were all calculated using alignments to the Kanost et al. assembly for reasons of gene model quality described below. Metrics related to pN/pS relied on SNP calls from the vcf files but estimates of π and Tajima’s D were performed directly from alignments using angd’s “saf2theta” command. In addition, we calculated ρ^2^ linkage disequilibrium for non-overlapping 50-basepair windows across the more contiguous Gershman assembly using the vcftools utility “--geno-r2”. These results are reported in Table 3 but we refrain from formal statistical comparisons between populations as the uneven sampling effort between them makes estimates of variation difficult to compare directly.

### Genetic characterization of differentiated regions

For each genomic region-of-interest, annotated genes in the region were examined based on the original assembly; this version’s annotations have a high degree of manual curation not present in the newer assembly (Kanost *et al*. 2016). *Manduca sexta* has a wealth of available RNA sequence data from both sexes and different life-stages, aggregated as a set of RNAseq collected by Cao and Jiang (2017) and quantified with this older reference. From this dataset, the specificity of gene expression was calculated for each gene in the genome for life-stage, tissue, and sex-specificity (following methods in Mongue, Hansen, & Walters, 2020) to inform potential function of genes-of-interest. More precisely, the specificity metric (SPM) was used to assess the proportion of a given gene’s expression found in a specific class. This metric ranges from zero to one, inclusive, and denotes the proportion of a given gene’s expression in a focal category (Kryuchkova-Mostacci and Robinson-Rechavi 2017). SPM was calculated for each of the three dimensions of specificity. Stage-specificity was defined as gene expression unique to larva, pupa, or adult *M. sexta* based on a set of tissues collected in all three stages (head, midgut, and fat bodies). Tissue-specificity within a stage was based on tissue types available for that stage (see Table 2). And, naturally, sex-specificity was based on amount of expression unique to males or females. For instance, a gene with an adult SPM of 0.90 shows 90% of its expression in adults as opposed to pupae or larvae. The same gene has separate SPM values that denote how specific its expression is to males or females and which tissues of those individuals. This method has the benefit of generating a sense of gene-function independent of functional annotation based on sequence homology with existing annotations.

**Table 2.**
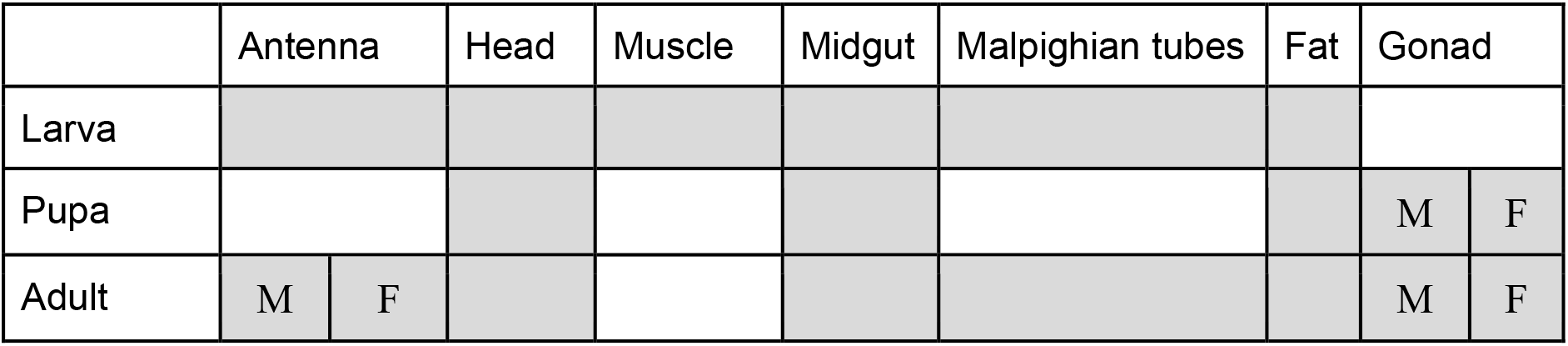
Tissue-specific RNA sequencing available for each life-stage of *M. sexta*. This dataset forms the basis for tissue specificity labeling annotation of genes in this study. Grey cells indicate presence, empty cells denote absence of data. If sex specific data are available, it is noted with an M (for male) and F (female) in the relevant cell. For larvae, few tissues were sampled at multiple instars, so no further developmental division was made.

**Table 3.** Population genetic parameters across the autosomes of the populations surveyed here. The Z chromosome was omitted due to uneven sampling of sexes between populations. Median values are given for polymorphism estimates (to avoid skew from outliers), while means are reported for Tajima’s D (as in every case, the median value is centered on zero). For all statistics involving silent and replacement sites (i.e. everything but linkage) the older Kanost et al. assembly was used to take advantage of higher confidence gene models. Mean linkage disequilibrium (ρ^2^) reported for 50 basepair windows using the Gershman et al. assembly to take advantage of more robustly supported linkage groups. Standard deviations appear in parentheses.

|  | Population genetic summary statistics for each population surveyed |  |  |
| --- | --- | --- | --- |
|  | North Carolina | Kansas | Arizona |
| pN | 0.0068<br>($\pm 0.022$) | 0.0044<br>($\pm 0.0198$) | 0.0063<br>($\pm 0.027$) |
| pS | 0.0232<br>($\pm 0.050$) | 0.0140<br>($\pm 0.050$) | 0.0197<br>($\pm 0.058$) |
| pN/pS | 0.2850<br>($\pm 0.656$) | 0.2852<br>($\pm 0.665$) | 0.2851<br>($\pm 0.652$) |
| $\pi_0$ | $8.82 \times 10^{-9}$<br>($\pm 0.030$) | $4.27 \times 10^{-8}$<br>($\pm 0.034$) | $7.95 \times 10^{-8}$<br>($\pm 0.032$) |
| $\pi_4$ | $5.96 \times 10^{-8}$<br>($\pm 0.082$) | $3.37 \times 10^{-7}$<br>($\pm 0.090$) | $4.32 \times 10^{-7}$<br>($\pm 0.086$) |
| Tajima's $D_0$ | -0.0795<br>($\pm 0.433$) | -0.0367<br>($\pm 0.299$) | -0.0673<br>($\pm 0.354$) |
| Tajima's $D_4$ | -0.0259<br>($\pm 0.510$) | -0.0148<br>($\pm 0.368$) | -0.0259<br>($\pm 0.406$) |
| $\rho^2$ | 0.3538<br>($\pm 0.396$) | 0.5990<br>($\pm 0.392$) | 0.4053<br>($\pm 0.401$) |

All genes across the genome were annotated with SPM values; however, for large outlier regions spanning numerous genes, it was not feasible to examine individual genes in depth. Instead, using the above annotations, these regions were examined for biased composition compared to non-differentiated regions. In other words, enrichment or depletion of functional classes within putative inversions was tested. For small regions containing one or a few genes, each could be manually examined. In these cases, single nucleotide polymorphisms in and around these genes were examined for evidence of coding-sequence variation between populations. SNPs were annotated with the tool SNPeff v. 4.2, which uses gene models to predict the coding sequence changes introduced by polymorphisms (e.g. loss of a canonical stop codon, Cingolani et al., 2012).

### Data accessibility

All newly generated sequencing data are available with the following accessions: SRP144217, PRJNA639154 on NCBI. Accessions for RNAseq used in functional inference can be found in (Cao and Jiang 2017). Custom scripts for analysis can be found at github.com/amongue/Manduca_Demography.git. New assembly chromosome assignments can be found in supplementary files.

## Results

### Chromosomal assignment of the new *M. sexta* assembly

The older, Illumina-only assembly of *M. sexta* already possesses chromosomal assignment based on syntenic alignments to *Bombyx mori*, a well-studied moth in a sister family to *M. sexta*. It has previously been shown that these two species share not only a conserved karyotype (n = 28) but also extensive conservation of sequence along these chromosomes (Yasukochi *et al*. 2009). Initially, we compared synteny between the two assemblies of *M. sexta* to maximize similarity and thus alignment information. From these alignments, 25 of the 28 largest scaffolds in the new assembly matched cleanly and overwhelmingly to one chromosome. The remaining three were more complicated and resolved as follows.

HiC_scaffold_31 showed synteny to both chromosome 2 and chromosome 26, in a way that suggested a potential fusion or misassembly, i.e. the first ~9Mb of the scaffold was syntenic to chromosome 26 and the remaining ~6Mb to chromosome 2. Closer inspection of published BAC-FISH mapping reveals that there has been structural turnover between species and both *B. mori* chr2 and chr26 are orthologous to a fused *M. sexta* chr2 (Yasukochi *et al*. 2009); thus this scaffold was correctly assembled and was manually annotated as chromosome 2.

Second, both HiC_scaffold_2 and HiC_scaffold_25 showed homology to *B. mori* chromosome 11 almost exclusively. Again, the BAC-FISH mapping revealed the reason for the anomaly. Chromosome 11 in *B. mori* is split in *M. sexta* into the larger chromosome 11 and smaller chromosome 26 (Yasukochi *et al*. 2009). Based on the relative sizes of these scaffolds, we annotated the larger (~10Mb) HiC_scaffold_2 as chromosome 11 and the smaller HiC_scaffold_25 (~5Mb) scaffold to chromosome 26.

With these manual annotations, each of the largest 28 scaffolds in the new assembly is now anchored to one of the 28 chromosomes in *M. sexta*. Smaller, more fragmented scaffolds were also assigned to chromosome based on syntenic alignment counts, but without manual curation. These assignments are presented in a supplementary file.

### Structure

We found three populations via the heuristic of Evanno et al. (2005): namely, the most probable K is the one the biggest change in likelihood from the previous K value. Per this metric, each of the three states’ moths form distinct populations, although there was some apparent admixture between Arizona and Kansas, including an Arizona individual confidently grouping with the Kansas samples (Figure 2a).

**Figure 2.**
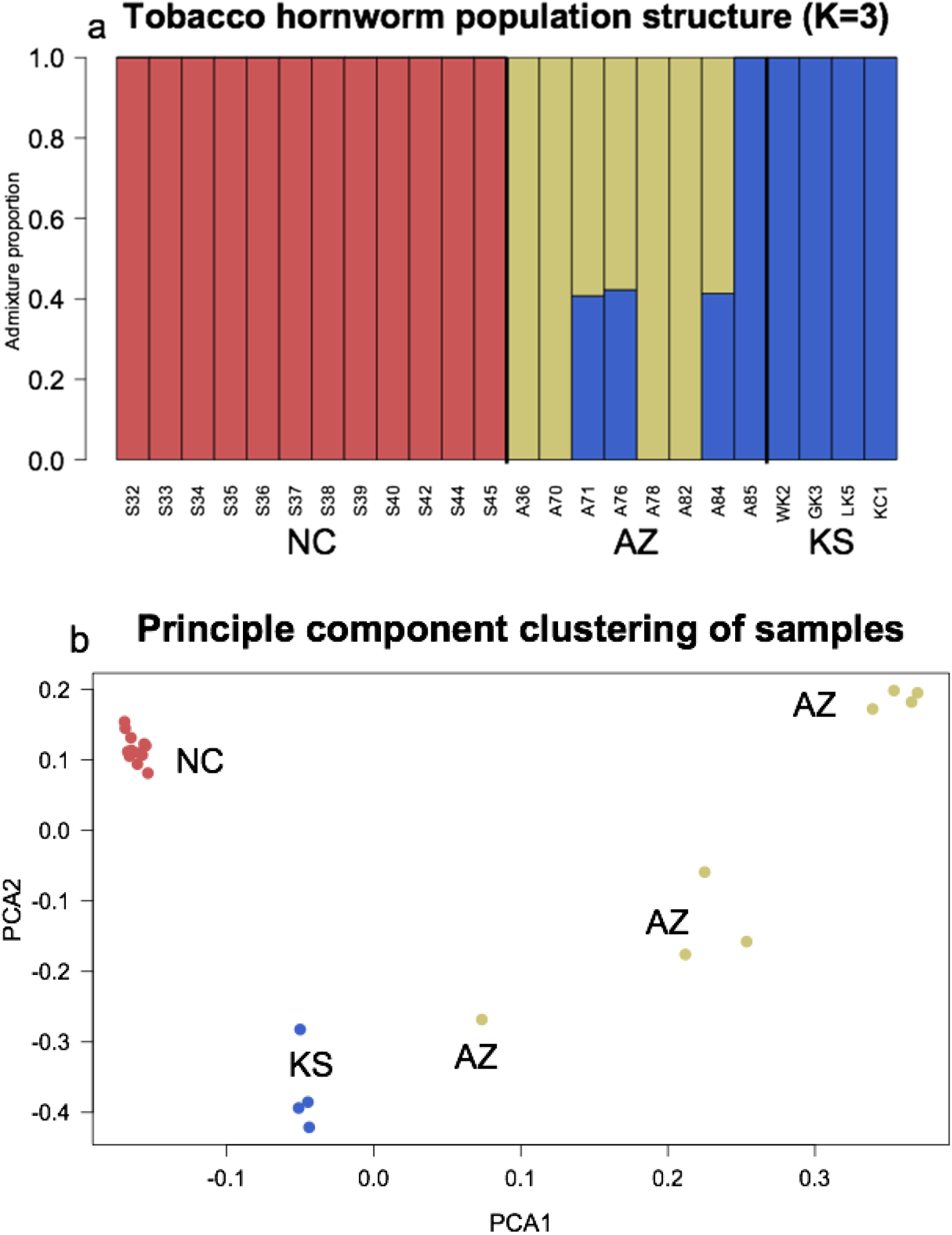
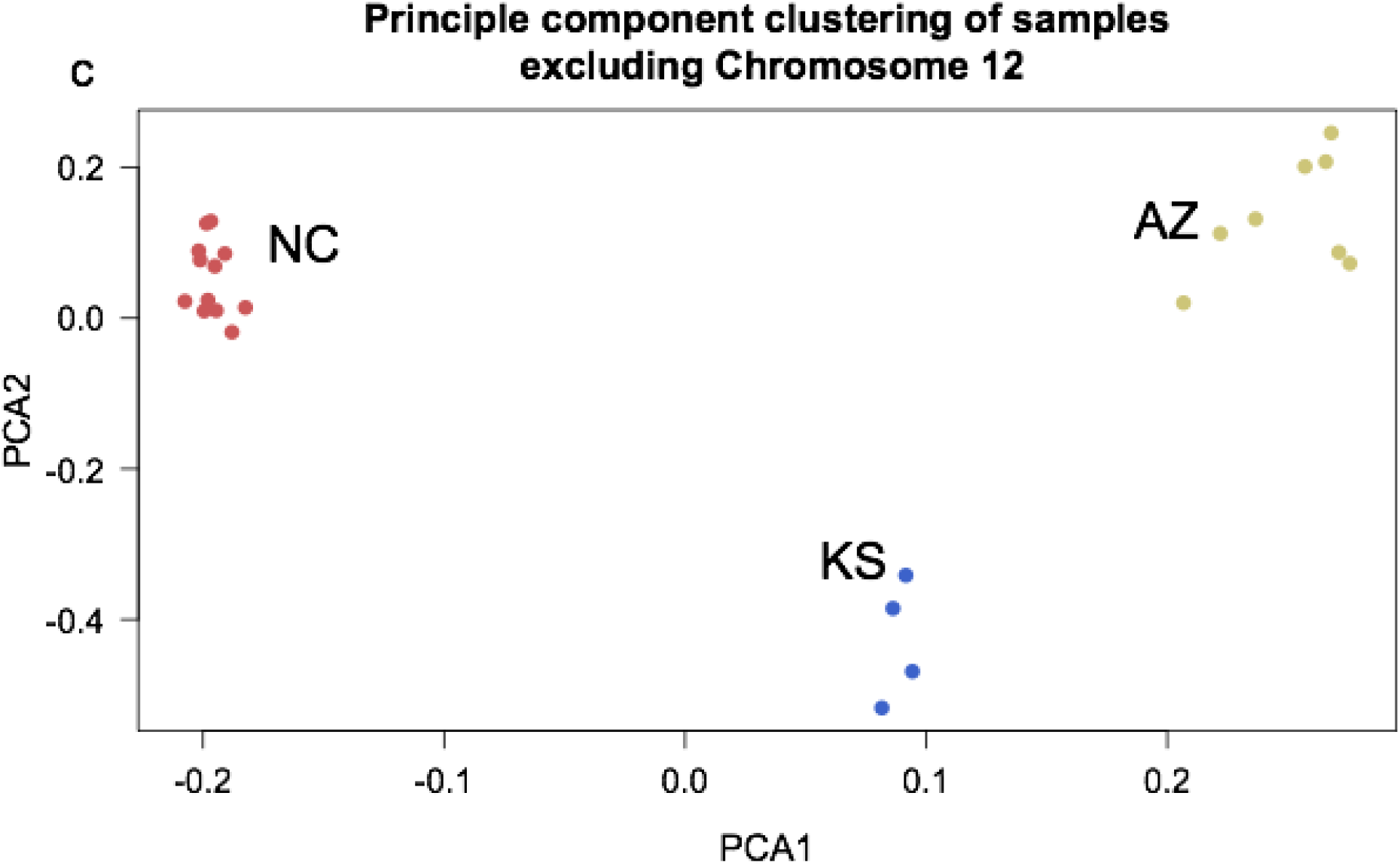
Population structure of *M. sexta* based on whole-genome resequencing of North Carolina (NC), Arizona (AZ), and Kansas (KS) individuals, omitting the Z chromosome. **(a)** Clustering of samples based on the most likely K (3), demonstrating that moths from different states are genetically differentiable. **(b)** Principle component analysis of the same dataset, with coloring the same as in (a). Note in both cases that Arizona individual (A85) clusters more closely with Kansas than other Arizona samples and three others (A71, A76, and A84) fall in between Kansas and the other four Arizona samples. Additional analysis revealed that these individuals lacked a large chromosome 12 inversion unique to the AZ population. **(c)** Excluding this chromosome from analysis causes all AZ individuals to cluster more closely with each other than with KS.

Taking a complimentary approach, we assessed principal components of SNP variation across our samples and observed the same clustering pattern as found in the structure analysis. Each state’s samples do not cluster with other states, but Arizona samples showing a more dispersed pattern than the others, including some individuals more similar to Kansas individuals than other Arizona individuals (Figure 2b). When excluding chromosome 12, the signal for admixture in some Arizonan moths disappears, and samples all sort by collecting location more cleanly (Figure 2c).

### Differentiation across the genome

To start, we provide population genetic summary statistics for each state in Table 3. Due to the unequal sampling effort between states, we do not formally test for statistical differences between populations. And indeed, absolute values for many measures of variation (e.g. pS or π) are lowest in Kansas, the population with the fewest sampled individuals. Still, the relative amount of non-synonymous variation scaled by synonymous variation (pN/pS) is remarkably consistent across populations, suggesting the strength of selection relative to drift is comparable across states. Moreover, Tajima’s D is more negative at zero-fold sites (i.e. those exposed to selection) than four-fold (putatively neutral sites) in each population, which further implies a role for directional selection in each population. Finally, apparent linkage disequilibrium is inversely correlated with sample size, as expected if artificially tight associations are observed between common alleles due to shallower sampling failing to capture rare alleles.

A simple genome-wide average of F_ST_ set a baseline level of differentiation of 0.04 to 0.09, depending on the pairwise comparison between states (Table 4). A sliding window scan across the genome revealed a few regions with markedly higher F_ST_ (> 0.5, Figure 3). From this analysis, several regions-of-interest became apparent. Most notably, a large proportion of chromosome 12 shows high levels of differentiation between Arizona and the other two populations. The Z chromosome showed the strongest peak of elevated differentiation, and chromosome 19 showed another above our threshold level. We compared these results on the newer assembly to a parallel F_ST_ scan on the older assembly (Figure S1). As expected, this turned up many differentiated scaffolds assigned to chromosome 12, the Z, and 19, but also 2, 6, and 13. In the following sections we describe each of these regions in detail and resolve apparent discrepancies.

**Table 4.** Genome-wide point estimates (excluding the Z chromosome) for differentiation between each pairwise combination of the three populations studied here. In each comparison differentiation at sites exposed to selection (i.e. zero-fold degenerate sites, 0x) and putatively neutral sites (four-fold degenerate sites, 4x) were estimated separately in angsd using the gene models from the Kanost et al. assembly.

| $F_{ST}$<br>0x, 4x | NC | KS |
| --- | --- | --- |
| KS | 0.064, 0.054 |  |
| AZ | 0.090, 0.081 | 0.041, 0.044 |

**Figure 3.**
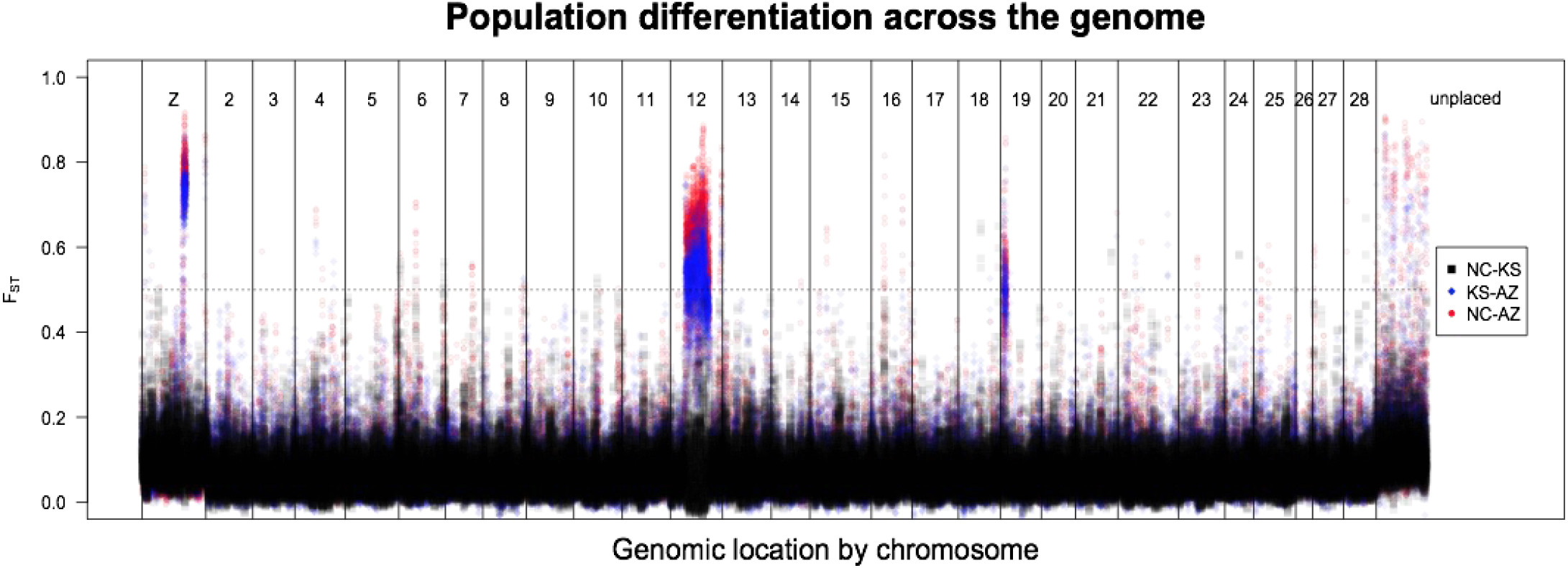
Pairwise differentiation (F_ST_) across the *M. sexta* genome: North Carolina (NC) vs. Kansas (KS; black squares), Kansas vs. Arizona (AZ; blue diamonds), and North Carolina vs. Arizona (red points). Plotting is ordered by chromosomal linkage, but not ordered within a chromosome. We examined regions enriched for differentiation greater than 0.5 (dashed line). By this metric, one peak on the Z, and one peak each on chromosomes 12 and 19. Additionally, note that the highest peaks almost always come from KS vs. AZ and NC vs. AZ, indicating that Arizona’s population is more distinct from the others than NC and KS are from each other. Chromosomes are numbered based on syntenic assignment of HiC scaffolds from the Gershman assembly. Unplace scaffolds are all those beyond the 28 main chromosomal scaffolds.

### An inferred segregating inversion on chromosome 12

We observed a large F_ST_ peak across a substantial portion of chromosome 12 in the more contiguous assembly (Figure 4). This peak corresponds with the Delly2 annotation of an inversion located on HiC_scaffold_12 from 5,414,046 Bp to 13,233,866bp, a total of 7.82Mb. Samples from North Carolina annotated with Delly lacked this called inversion, as expected given that the reference individual was sourced from lab stock which was originally collected in North Carolina.

**Figure 4.**
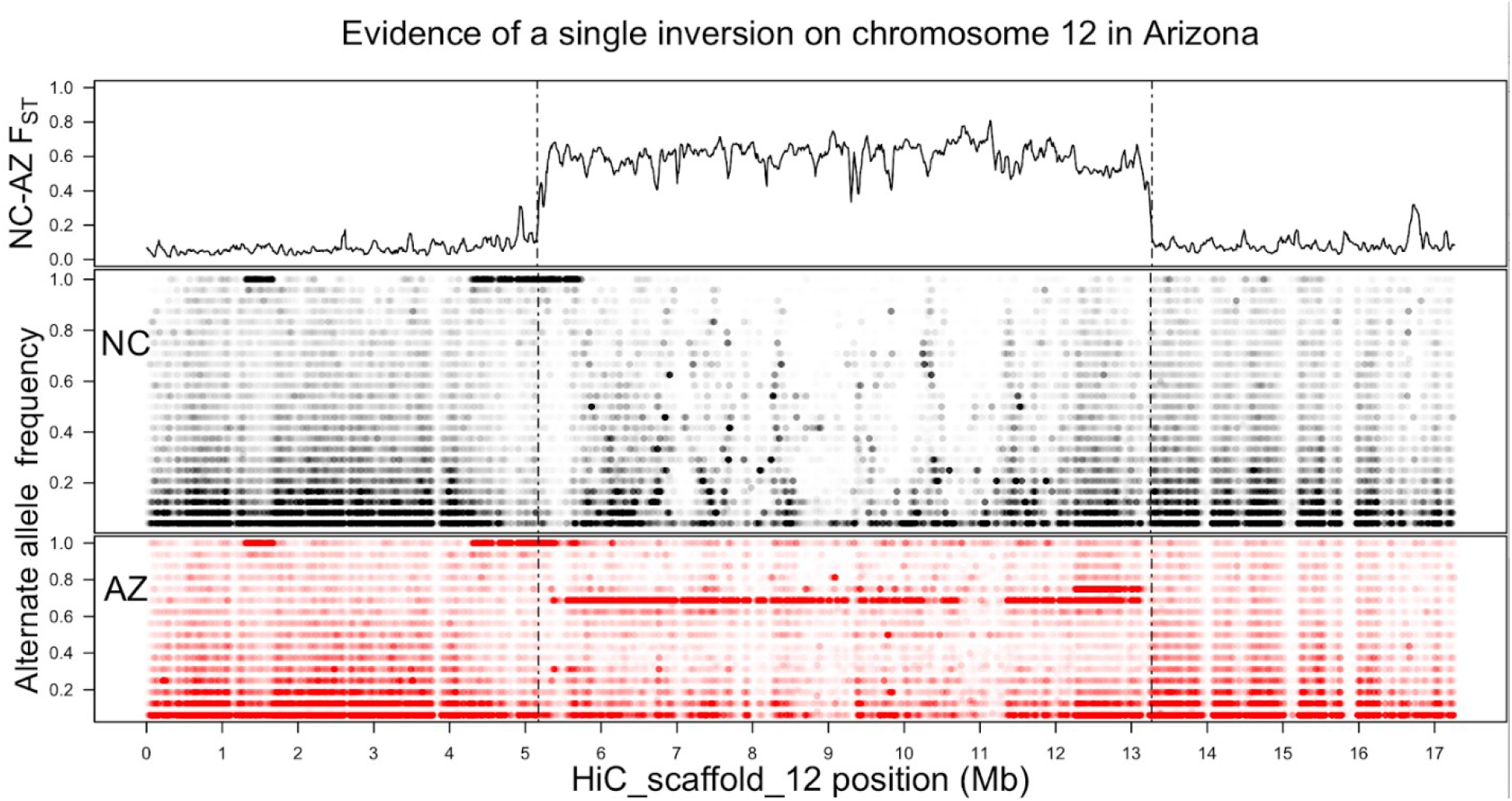
Evidence of a single segregating inversion on chromosome 12 from the newly released *M. sexta* assembly (Gershman *et al*. 2021). **Top.** F_ST_ comparison for the North Carolina and Arizona populations shows a roughly 8 Mb region (between 5.3-13.3 Mb) of elevated differentiation between the populations, corresponding to the length of the two tracts of differentiation from Figure 3, plus some additional unplaced peaks. Vertical dashed lines delimit this differentiated region and extend below for ease of comparison between populations. **Middle and Bottom.** Frequencies of alternate alleles for called SNPs in North Carolina (middle) and Arizona (bottom). Points are semi-transparent such that opaquer regions are denser with variants. Arizona appears to hold the inverted orientation as a single allele frequency (0.6875 or 11/16) dominates the differentiated region, whereas North Carolinian allele frequencies are variable throughout. Also in the 1-2 Mb and 4-5 Mb regions are areas of dense SNPs fixed in both populations. These are interpreted as regions in which the lab strain used to generate the reference has differentiated from the wild populations.

In the old assembly, this peak correspond to 13 scaffolds, listed in Table S2 and plotted in Figure S2, summing to 7.6 Mb of sequence, approximately 56% of the total (assigned) length of the chromosome. The peaks contain 321 annotated genes. Note the discrepancy of ~200Kb of differentiated sequence.

The putative inversion’s frequency of 0.6875 corresponds to 11 of 16 sampled chromosomes with the inversion. Per investigation of sample genotypes, this variation is distributed as 4 individuals homozygous for the non-reference orientation (A36, A70, A78, A82), three heterozygous for it (A71, A76, A84), and one homozygous for the ancestral orientation (A85). Note that the heterozygous individuals correspond to those inferred to be admixed with Kansas and the one homozygous non-inverted individual was grouped with Kansas completely in structure analyses. As discussed above, removing chromosome 12 from structuring analyses leads to less similarity between AZ and KS samples, suggesting the inversion presence/absence is large portion of the signal of the Arizona population. A scan of linkage disequilibrium along the chromosome (Figure S3) showed elevated linkage within the bounds of the hypothesized inversion in each of the three populations, suggesting that this region is a barrier to geneflow between populations.

The large size of this inferred inversion (roughly 8Mb and 321 genes contained within) makes identifying a causal adaptive locus challenging. We used the RNA sequencing metadataset to classify genes as specific to one sex (male or female), life stage (larva, pupa, or adult), and tissue (antennae, head, fatbody, midgut, Malpighian tube, muscle, ovaries, or testes) if a given gene showed 70% or more of its expression exclusively in that category. By this metric, the putative inversion on chromosome 12 does not differ in sex-biased composition (X^2^_2_ = 1.31, p = 0.518), lifestage-biased composition (X^2^_3_ = 2.06, p = 0.560), or tissue-biased composition (X^2^_7_ = 7.45, p = 0.384) compared to the rest of the genome. Thus, even if there is a set of locally adaptive alleles in this region, it is not one enriched for any of the annotations available to us. Nevertheless, we identified one gene of interest in this region thanks to a misassembly.

Both the ostensible chromosome 2 and 13 peaks from the more fragmented assembly belong to the putatively inverted region of chromosome 12 in the newer assembly. The former is rather unremarkable, some 50Kb of a scaffold containing two annotated genes (Msex2.02572 and Msex2.02573) with highly differentiated SNPs localized to areas outside the coding region of either. These variants may be truly neutral or impact the regulation of the nearby genes, but without a better understanding of regulatory elements (e.g. promoter sequences) in this species, it is impossible to say. The peak previously assigned to chromosome 13, deserves more consideration however.

### A potential pseudogene within the Chromosome 12 inversion

Despite overall congruence of results from the old and new assembly, one difference in the F_ST_ scans proved particularly interesting. The older assembly indicated a peak of differentiation on a scaffold anchored to chromosome 13; however, none of the more contiguous chromosome 13 is particularly differentiated using the new assembly as a reference. To reconcile this contradictory pair of results, we investigated the peak in the old assembly first, then placed it in new assembly.

This peak comes from one scaffold (scaffold00381) starting around 200Kb and continuing for the remaining 100Kb (Figure 5a). This region contains a single gene, Msex2.10493, and it displays a striking pattern of differentiation between states. Within population variation is low in Kansas and North Carolina, with only 1 and 3 coding-sequence polymorphisms observed, respectively; in contrast, 61 SNPs were called in the gene body from the Arizona samples (summarized in Figure 5b). Moreover, many of these SNPs are at high frequency (Figure 5c), including the predicted loss of a stop codon.

**Figure 5.**
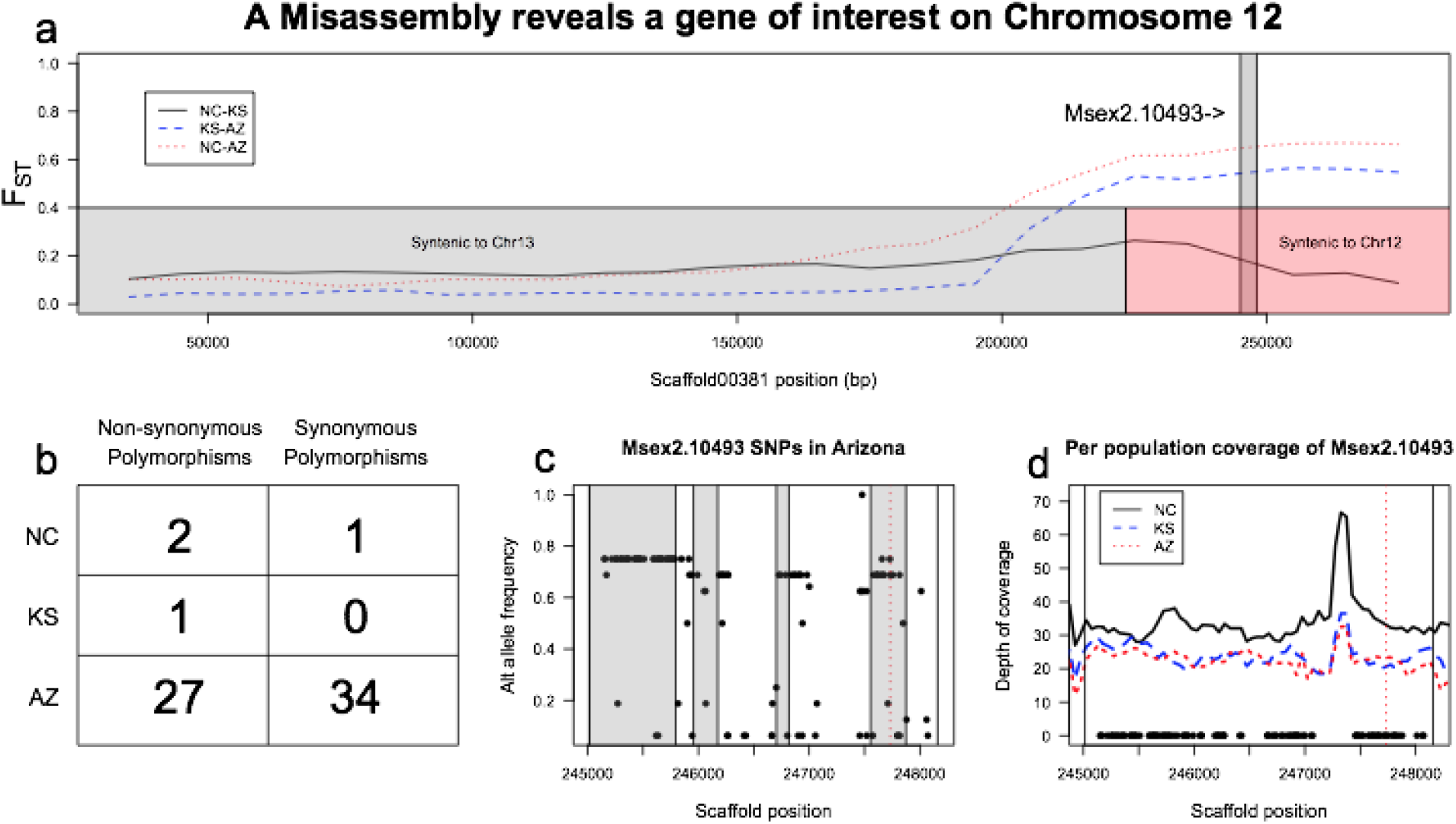
Extreme population differentiation of a gene within chromosome 12’s putative inversion. **a.** Finer scale plotting of F_ST_ of the differentiated Kanost et al. scaffold, previously assigned to chromosome 13 shows that differentiation occurs beyond 200 Kb and, as with the inversions above, it appears that Arizona is markedly different from North Carolina and Kansas. Boxes at the bottom of the plot show syntenic alignments with the Gershman et al. assemble and reveal that the differentiated region belongs to chromosome 12 rather than 12. A single annotated gene, Msex2.10493, lies in this region (grey box). **b.** Annotation of SNPs in the coding region reveals a stark difference in variation between populations. Arizona moths hold more than ten times the non-synonymous polymorphisms than those in the other two states. **c.** Frequencies of the SNPs in Arizona. Grey boxes represent exons and the dashed vertical line marks a variant that disrupts the annotated stop codon. **d.** Depth of coverage of the gene in question does not vary between populations in a way that suggests differences in alignment between populations can explain the differences in called SNPs.

The massive difference in called variants theoretically could arise from population-specific differences in mapping rates in this region, either through poor mapping in populations with low SNP counts or (far less likely) a pile-up of reads in Arizona owing to a collapsed segregating paralog. To rule out these artefactual explanations, we examined per-base coverage around this gene and found no differences between populations (Figure 5d; note that slightly deeper coverage in NC is consistent with genome-wide differences in sequencing effort between the populations). Thus, Arizona *M. sexta* appear to truly differ from the other populations with regard to the molecular evolution of this gene, which begs the question of the gene’s functional role. Msex2.10493 is overwhelmingly (98.2%) expressed in larvae compared to 1.8% adult expression and vanishingly small pupal expression. Within larvae, this gene shows 97% of its expression in the antennae and another 2.8% in the head.

The final question to consider was why such a striking pattern of difference is absent in the newer assembly. To answer this, we examined the syntenic alignment of scaffold00381 to the more contiguous assembly. A discrepancy immediately arose: roughly 2/3^rds^ of the scaffold aligned to HiC_scaffold30 (which we have anchored as chromosome 13), but the remaining 1/3^rd^, including the coding region for Msex2.10493, aligns to chromosome 12 (Supplemental Figure S4). This pattern is suggestive of a misassembly in the older reference genome. Furthermore, the roughly 100Kb length of this apparently chromosome 12 sequence accounts for half of the 200Kb discrepancy in the estimated chromosome 12 differentiation peak between the two assemblies.

More direct evidence suggests that this potential pseudogene lies within the chromosome 12 inversion. First, the coding sequence of the old assembly’s Msex2.10493 is homologous to the new assembly’s XM_037442453.1, which lies in the inverted region (roughly 9.257 - 9.261Mb into chromosome 12). Additionally, the frequency of the stop-loss is 0.6875 ( = 11/16) and matches that of the most common SNP frequency within the inversion. Moreover, the genotypes (homozygous present, heterozygous, homozygous absent) of the inversion perfectly match the stop-loss variant’s genotype. In conclusion, this potential pseudogene sits in the middle of the inferred inversion on chromosome 12, but it was only identified thanks to a serendipitous misassembly in the older version of the *M. sexta* reference.

### The Z holds another putative segregating inversion

The Z chromosome’s outliers are attributable to roughly 1Mb in the middle of the chromosome-level scaffold (HiC_scaffold_19). The scale of elevated differentiation suggests another inversion, and our structural variant scan identified an inversion that precisely overlaps this region (from 13,706,400 to 14,710,588 bp, for a total length of 1,004,188 bp). Further evidence for a lack of recombination in this region can be seen in the non-reference allele frequencies which vary across the scaffold in both NC and AZ but show a tract of SNPs with a common frequency of 0.93 (= 13/14) within the putative inverted region in Arizona (Figure 6).

**Figure 6.**
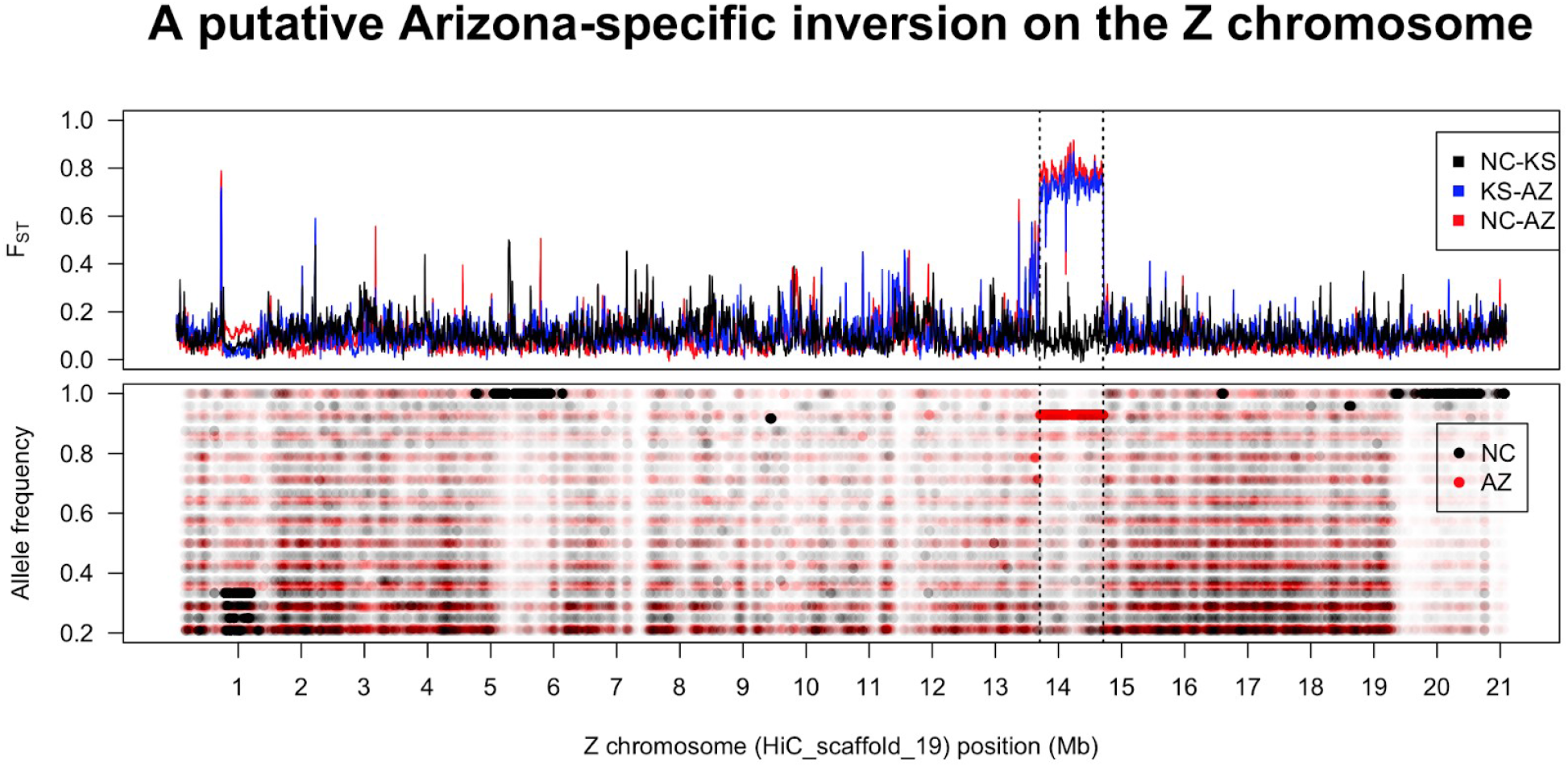
Evidence for an inversion on the Z chromosome. **Top.** A roughly 1 Mb portion of the Z shows elevated differentiation in North Carolina – Arizona and Kansas – Arizona comparisons. Here, as well as below, this region corresponds to an inversion identified by Delly2 (from 13.7 to 14.7 Mb.). **Bottom.** Like the chromosome 12 inversion, Arizona shows a long tract of shared non-reference allele frequencies in the region while North Carolina does not (note that Kansas allele frequencies are omitted due to lower sampling effort artificially stratifying frequencies). Inferred blocks of non-reference SNPs that do not correspond to similar F_ST_ peaks are inferred to be regions in which the lab-strain used for sequencing has differentiated from the natural populations.

This peak of differentiation is also identifiable in the older assembly as a roughly 1Mb region of scaffold00022, suggesting we are capturing essentially all of the inversion on one scaffold, even in the more fragmented assembly, (Figure S5). In terms of gene content, scaffold00022 contains 35 genes and shows an significant excess of unbiased genes (i.e. those expressed in both sexes, X^2^_2_ = 11.36, p = 0.0034), compared to the rest of the Z, which is significantly masculinized, as previously reported (Mongue *et al*. 2020).

### Chromosomes 6 and 19 show differentiation but no clearly differentiated genes

Chromosome 6 shows a single peak of divergence, though this region is only weakly differentiated, barely meeting our cut-off threshold for exploration (F_ST_ > 0.5 between any two populations). This region contains three full annotated genes (Msex2.12148, Msex2.12149, and Msex2.12150), but no SNPs within coding regions.

Two outlier scaffolds in the older assembly localize to chromosome 19; both represent differentiation between Arizona and the other two states. In the newer, more contiguous assembly, these peaks remain distinct from each other (Figure S6), suggesting they are not parts of a common feature. One is localized to scaffold00118, to approximately the last 400Kb of an ~850Kb scaffold. This region contains 11 annotated genes (Msex2.05775-Msex2.05786), but none of these shows skews in coding region SNPs between populations as extreme as seen on the potential pseudogene mentioned above. However, two genes do show more modest deviation between states. Msex2.05777 is variable in both North Carolina (Pn = 6, Ps = 10) and Kansas (Pn = 7, Ps = 12) but monomorphic (Pn = 0, Ps = 0) in Arizona. Conversely, Msex2.05784 is non-variable in the North Carolina and Kansas samples but contains 3 non-synonymous variants and 1 synonymous variant in Arizona.

The second peak, on scaffold00419, seems to span the length of this 200Kb scaffold, but this may reflect the limits of resolution of our F_ST_ window-size. In any case, the entire scaffold contains 11 annotated genes, many of which have no expression data and no called SNPs in any population. As such, we do not place much emphasis on the differentiation around these dubiously annotated regions.

## Discussion

We conducted whole genome resequencing to investigate the natural variation of *Manduca sexta* populations across the United States. In brief, we found that individuals from three populations across an East-West transect are genetically differentiated. This result may not seem surprising, but studies of other Lepidoptera have at times found evidence for continent-wide panmixis (specifically for monarch butterflies, Lyons et al., 2012; Zhan et al., 2014). And while monarchs may be an extreme case of lepidopteran long-distance travel, sphinx moths, including *M. sexta*, are renowned for their flight and dispersal ability, both in terms of small-scale speed and dexterity (Stevenson *et al*. 1995) and long-distance dispersal (Janzen 1984; Haber and Frankie 1989). Indeed, *M. sexta* has been posited as migratory in the past (Ferguson 1991). Our results, however, suggest that gene flow is less common than expected of a species with routine migration. In fact, the genome-wide mean F_ST_ for the least differentiated pair (0.04, Kansas-Arizona) is on par with the differentiation between migratory North American monarchs and the Hawaiian population that is thought to have split roughly 200 years ago (Lyons *et al*. 2012).

More striking than this level of differentiation is the nature of the differences that contribute to it. Namely, large (>1 Mb) putative inversions, are key contributors to the variation between populations. Sustained tracts of differentiation are most apparent on two chromosomes, the Z chromosome and chromosome 12. In both cases, the Arizona population is distinct from Kansas and North Carolina. We note that the smaller sample size of Kansas *M. sexta* decreases the precision of allele frequency estimation (Lou *et al*. 2020), and may obscure some of the more subtle population differences for the time being. For the present study, we discuss in detail the genomic and genetic nature of the large differences, with the caveat that functional genetics will be required to truly confirm the genomic structure and downstream fitness consequences of this variation.

### The largest signal for population differentiation: a likely inversion on chromosome 12

A long tract of chromosome 12 showed high levels of differentiation in our F_ST_ scan. This signal belonged to a single block of sequence, spanning 8 megabases (roughly half the length of the chromosome), containing over 300 annotated genes, and found at a frequency of almost 70% in the Arizona population. Individuals that are heterozygous for this putative inversion or lacking it entirely cluster more closely with the Kansas population than other Arizona samples which are homozygous for it. When removing chromosome 12 from analyses, all Arizona samples cluster more closely with each other than with Kansas however, suggesting that the two populations are differentiated across the genome, but that the sheer number of variants associated with the putative inversion swamps other signals. These two observations combine to suggest that the inversion has been segregating in the population for some time.

In general, and especially in large populations, inversion frequencies are thought to imply something about their selective effects, with only beneficial variants reaching high frequency (Lande 1984; Kirkpatrick and Barton 2006). In other species in which naturally occurring inversions have been studied, this expectation has been borne out. Low frequency inversions (e.g. 8%) are associated with fitness tradeoffs (Lee *et al*. 2016) and more common inversions (~50%) may have little to no fitness costs (Knief *et al*. 2016), at least locally. Their absence elsewhere may suggest a lack of gene flow or that there *are* fitness costs in other environmental backgrounds. Indeed, linkage disequilibrium is higher within the bounds of the inverted region even in populations in which we did not detect the variant, suggesting a dampening of gene flow between populations here. From this perspective, the common chromosome 12 variant seems likely to be at least selectively neutral, if not beneficial, in Arizona. However, the large number of encompassed genes present a challenge for identifying selectively relevant loci from purely computational methods. Still, one gene-of-interest arose from our efforts to cross-check results with the older *M. sexta* reference (Kanost *et al*. 2016).

### A likely candidate for local adaptation in the inversion: an apparent pseudogene

An F_ST_ scan of the older assembly with the same resequencing data revealed numerous outlier scaffolds anchored to chromosome 12 as expected. However, an additional similar peak was detected on a scaffold assigned to chromosome 13. Alternate allele frequencies on this peak matched those most commonly seen in the chromosome 12 inversion, and a closer inspection of synteny between the old and new assemblies suggested that this scaffold is likely a chimera. The majority of sequence belongs to chromosome 13, but the differentiated region is syntenic to chromosome 12 within the inversion. This region contains a single annotated gene, Msex2.10493 (in the old annotation; XM_037442453.1 in the new annotation). It carries almost no variation in North Carolina or Kansas but has a wealth of high frequency variants (61 SNPs in our sample) in the Arizona population, one of which is a predicted stop-loss in the coding region. A large-effect SNP like this could easily break the protein function via translation of otherwise non-coding DNA. While inconclusive from molecular data alone, a hypothetical loss of function would easily explain the preponderance of SNPs found in Arizona; pseudogenization instantly relaxes selective pressure such that subsequent variants can accrue quickly. And indeed, the high frequency of this stop-loss suggests that it cannot have an appreciable fitness cost, or it would not have persisted in the population. Naturally, this raises the question of what function this gene has that it could be lost without strong selective consequences. Or, alternatively, could a loss of function be adaptive under certain circumstances?

Based on our expression characterization, the Msex2.10493 gene is almost exclusively expressed in larval antennae, which are known to play a role in olfaction and food choice in *M. sexta* (Boer and Hanson 1987). Olfaction genes are among the most commonly observed to pseudogenize in other species, ostensibly because the fitness consequences of their loss are rarely lethal (Nei *et al*. 2008; Prieto-Godino *et al*. 2016). Intriguingly, while most populations of *M. sexta* feed almost exclusively on plants in the family Solanaceae, larval Arizona hornworms have been recorded eating the unrelated devil’s claw (Martyniaceae: *Proboscidea parviflora*, Mechaber & Hildebrand, 2000). It is tempting to make the connection between the apparent loss of function of a larval sensory gene and the exploitation of a new hostplant, as research on the silkmoth (*B. mori*) has shown that knockout of a single chemoreceptor is sufficient to turn naturally specialist larvae into generalists (Zhang *et al*. 2019); however, it will take further study to establish whether or not a similar situation is occurring in *M. sexta*.

Still, the placement of this apparent pseudogene within a massive block of elevated differentiation between populations and its potential relationship with hostplant ecology is suggestive of a locally adapting region. As genomic scans of natural variation become more common, so too does the observation that many species harbor regions of large-scale differentiation that distinguish individuals adapted to different environments (Fuller *et al*. 2019; Todesco *et al*. 2020; Mérot *et al*. 2021, 2021). In such instances, suppressed recombination, typically through inversions, keep locally beneficial alleles from being decoupled from each other, even in the face of gene flow from other populations. The frequencies of these large variants typically follow an environmental (or more directly a selective) cline (Kapun *et al*. 2016; Wellenreuther *et al*. 2017). We do not have the geographic resolution to detect any potential cline in the frequency of this inversion and pseudogene, but presence and absence data are at least consistent with a role in expanded hostplant usage.

The devil’s claw plant’s native range is limited to the desert southwest of North America (particularly: northern Mexico and California, Arizona, New Mexico, Utah, and Nevada in the United States; see Berry *et al*. 1981). As expected based on this range, we did not observe any *M. sexta* carrying the putative inversion in North Carolina or Kansas. With only four sampled individuals from Kansas, we cannot exclude the possibility that the Arizona inversion is present there as well, albeit at a much lower frequency, but to follow the logic of the devil’s claw hypothesis, there would be no local benefit for individuals carrying the pseudogene outside of the southwest. Indeed, in the absence of a novel hostplant with the correct set of complex phytochemistry, becoming less choosy could be maladaptive. Zhang et al. (2019) found that although chemoreceptor-knockout silkworms readily ate many food sources, they failed to finish development and pupation on all but the traditional hostplant. To reiterate, these inferences are highly speculative, but they create easily testable hypotheses. Laboratory studies of Arizonan *M. sexta* with the putative inversion and pseudogene will be needed to unpack the functional consequences of this striking variation.

### A Z inversion in AZ: a role for sex-bias driving rearrangement

The single highest peak of differentiation in our genome-wide scan comes from the Z chromosome. This region, with roughly 1 Mb of extended differentiation and shared allele frequencies in Arizona also fits the pattern of a segregating inversion between populations. Moreover, the non-reference orientation is nearly fixed in the Southwest population, being homozygous (or present in hemizygous females) in all but one individual. As discussed above, an inversion is unlikely to reach such high frequency without some selective advantage.

Here, even without specific gene functional annotations, we have a reasonable hypothesis for what could drive this structural variant to near-fixation. We found that the inversion contains a significant excess of genes expressed in both sexes (i.e. unbiased genes), compared to the rest of the Z chromosome, which is male-biased (Mongue and Walters 2017). As has been recently shown, unbiased genes are an important source of positive selection on the Z in *M. sexta* (Mongue *et al*. 2021). These genes are expressed in a haploid state in females (which have only one Z chromosome); thus, even recessive beneficial mutations are exposed to positive selection (Rice 1984; Charlesworth *et al*. 1987). The capture of positively selected variants of such genes in a non-recombining region would ensure their co-transmission and seems a reasonable explanation for this inversion’s prevalence.

Moreover, these sex chromosome dynamics intersect with the population genetics of local adaptation in synergistic ways. Modeling suggests that the exposure of recessive alleles to selection should facilitate local adaptation and make the sex chromosomes hotspots for this phenomenon (Lasne *et al*. 2017). For similar reasons, sex-linked inversions are thought to sweep to fixation more easily than autosomal inversions (Connallon *et al*. 2018). As such, it is unsurprising to find the Z involved in short-term differentiation as well as long term adaptation (Mongue *et al*. 2021) in this species.

### Other differentiated regions of the genome

The inversions discussed above are the most apparent sources of regional differentiation, but chromosomes 6 and 19 also showed elevations in differentiation at smaller scales. For the most part, differentiated regions fell outside of coding regions or near dubiously annotated genes (i.e. those not supported by expression data). Lacking further information for hypothesized importance, they are not discussed in detail here for brevity’s sake.

## Conclusions

We report that *M. sexta* from Arizona, Kansas, and North Carolina are genetically differentiated. The confirmation of regional differentiation is a valuable addition to research in both laboratory and field studies, as well as to development of management plans for this crop pest between regions. That said, we have yet to establish the minimum geographic scale at which *M. sexta* populations are structured (for instance how much gene flow exists between adjacent states or even within a single state). This finer scale sampling will be particularly important for agricultural centers on the applied side, and additional sampling will also provide more insight into the large-scale variation we report here.

At least two large inversions and a potential pseudogene differentiate Arizona from the Kansas and North Carolina populations. All of these variants are high frequency (> 0.5) within Arizona but absent in the other populations. However, the nature of our sample collection limits the inferences we can make at present. For instance, the high frequency of the variants suggests they are not deleterious, but with only a single time-point of sampling in Arizona, it is hard to say whether or not these variants have reached an equilibrium frequency. Additionally, other studies have shown inversion polymorphisms to follow geographic or environmental clines (Anderson *et al*. 2005; Wellenreuther *et al*. 2017). In such cases, the inversions often hold locally adaptive alleles that are maintained in spatial polymorphism by differences in environmental factors, but for *M. sexta*, with each sampled state separated from the others by at least 1,500 kilometers, it is impossible to say if these segregating variants also follow a cline in the region or to which climactic variables they may relate. Additional sampling of the Southwest will be required to assess how widespread the inversions are.

Likewise, controlled laboratory studies will be required to assess the fitness consequences of the segregating stop-loss in a larval antennal gene within the chromosome 12 inversion. In particular, the coincidence of a novel larval hostplant in Arizona and a large-effect mutation in a gene expressed in a larval chemosensory gene are suggestive but far from conclusive. The relationship between the two will require hostplant choice experiments for larvae with and without the stop-loss mutation.

Finally, it is worth noting that these dramatic differences between populations could not be observed in standard lab strains, which are all derived from the North Carolina population to our knowledge. On the other hand, without extensive lab studies and RNA-sequencing, we would not have the information to form functional hypotheses for these variants found in wild populations. Beyond the immediate utility of these data, the results we present here serve as a reminder of the efficacy of integrating laboratory and field research within the same study system.

## Acknowledgements

This project was partially funded by NSF DDIG to AJM (DEB-1701931) and DEB #1557007 and IOS #1920895 to AYK. The authors wish to acknowledge Wesley Mason and Michael Hulet and the rest of the Information and Telecommunication Technology Center (ITTC) staff at the University of Kansas for their support with our high-performance computing. The authors also thank the University of Kansas’s Entomology Endowment for additional funding support in sequencing. Thanks to Paul Hime for guidance on population structure analysis. Thank you to Clyde Sorenson, Judie Bronstein, and Goggy Davidowitz, for initial and continued support in investigating population genomics of *M. sexta*. Kelly Dexter and Rebecca Messcher helped provide the samples from University of Florida’s McGuire Center collection. Thanks also to the members of Ashworth 2.02 for continued patience in the discussion of the material of this manuscript.

## Supplementary tables and figures

**Table S1.** Scaffold information for genomic regions of interest. To be included in this table, a scaffold had to show at least one 10 kilobase window in which F_ST_ was greater than 0.5 between any two of the 3 populations. The **Type** of differentiation (e.g. localize peak vs. inversion) was determined through subsequent analyses. For convenience, genomic locations are provided for both assemblies of *M. sexta*.

| Putative chromosome | Kanost *et al.* Scaffold | Gershman et al. Scaffold | Type |
| --- | --- | --- | --- |
| Z | scaffold00022 | HiC_scaffold_19 | Inversion |
| 12 | scaffold00042 | HiC_scaffold_12 | Inversion |
| 12 | scaffold00095 | HiC_scaffold_12 | Inversion |
| 12 | scaffold00119 | HiC_scaffold_12 | Inversion |
| 12 | scaffold00158 | HiC_scaffold_12 | Inversion |
| 12 | scaffold00203 | HiC_scaffold_12 | Inversion |
| 12 | scaffold00240 | HiC_scaffold_12 | Inversion |
| 12 | scaffold00264 | HiC_scaffold_12 | Inversion |
| 12 | scaffold00300 | HiC_scaffold_12 | Inversion |
| 12 | scaffold00335 | HiC_scaffold_12 | Inversion |
| 12 | scaffold00460 | HiC_scaffold_12 | Inversion |
| 12 | scaffold00466 | HiC_scaffold_12 | Inversion |
| 12 | scaffold00571 | HiC_scaffold_12 | Inversion |
| 12 | scaffold00616 | HiC_scaffold_12 | Inversion |
| 12 | scaffold00633 | HiC_scaffold_12 | Inversion |
| 12 | scaffold00790 | HiC_scaffold_12 | Inversion |
| 12 | scaffold00888 | HiC_scaffold_12 | Inversion |
| 12 | scaffold00921 | HiC_scaffold_12 | Inversion |
| 12 | scaffold00966 | HiC_scaffold_12 | Inversion |
| 12 | scaffold01138 | HiC_scaffold_12 | Inversion |
| 12 | scaffold01198 | HiC_scaffold_12 | Inversion |
| 12 | scaffold01282 | HiC_scaffold_12 | Inversion |
| 12 | scaffold01317 | HiC_scaffold_12 | Inversion |
| 12 | scaffold01366 | HiC_scaffold_12 | Inversion |
| 12 | scaffold01482 | HiC_scaffold_12 | Inversion |
| 12 | scaffold01552 | HiC_scaffold_12 | Inversion |
| 12 | scaffold00381 | HiC_scaffold_12 | Inversion,<br>potential pseudogene |
| 19 | scaffold00118 | HiC_scaffold_21 | Localized peak |
| 19 | Scaffold00419 | HiC_scaffold_21 | Localized peak |

**Table S2.**
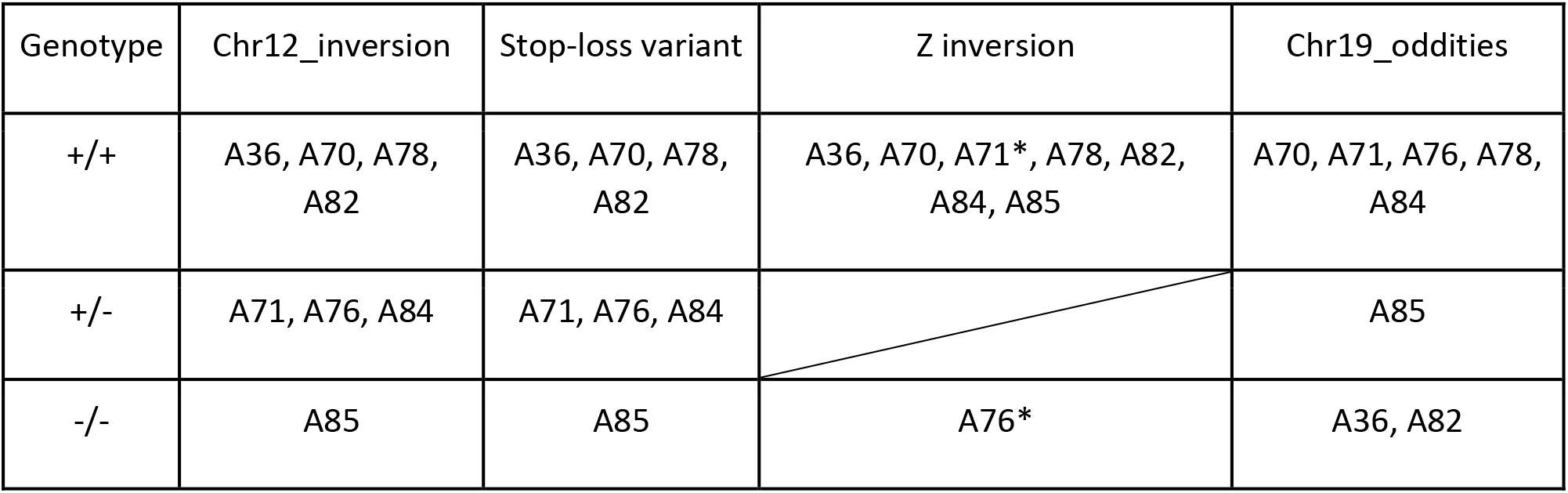
Genotypes of Arizona individuals for each of the key variants discussed in this manuscript. + denotes presence, - absence. The stop-loss variant and chromosome 12 inversion co-segregate in perfect linkage, further proof that the former resides within the latter. The Z inversion is not structured in the same way, however. Individuals marked with an * are female and hemizygous. Thus A71’s single Z carries the inversion while A76’s lacks it.

**Figure S1.**
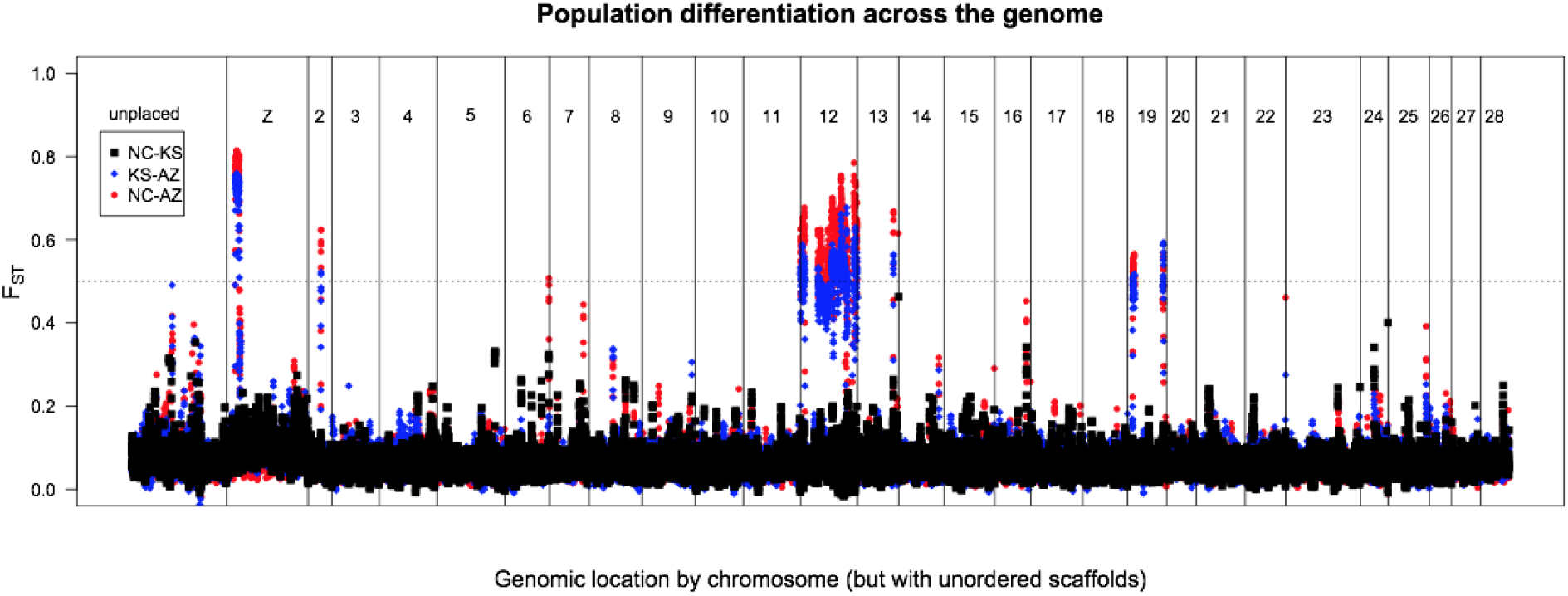
Pairwise differentiation (F_ST_) across the *M. sexta* genome using the Kanost *et al*. reference: North Carolina (NC) vs. Kansas (KS; black squares), Kansas vs. Arizona (AZ; blue diamonds), and North Carolina vs. Arizona (red points). Plotting is ordered by chromosomal linkage, but not ordered within a chromosome. We examined regions of differentiation greater than 0.5 (dashed line). By this metric, one peak on the Z, and one peak each on chromosomes 2, 13, and two on 19. Additionally, note the massive spike on chromosome 12, made up of numerous highly differentiated scaffolds. Some assignments, especially the peak on chromosome 13, were updated in the new assembly.

**Figure S2.**
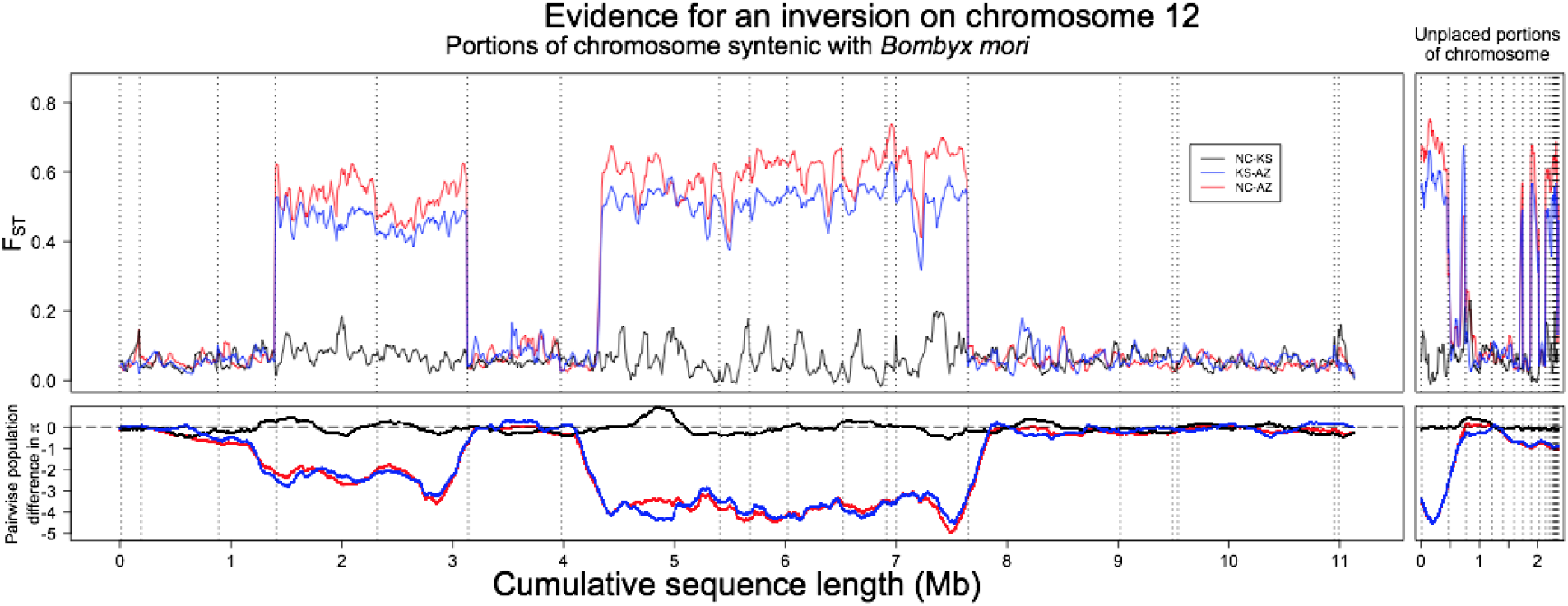
Investigation of differentiation across chromosome 12. **Top:** Ordering scaffolds from **Figure S1** via synteny with *Bombyx mori* suggests two regions > 1.5 Mb of elevated differentiation (F_ST_) between *M. sexta* populations. Given the wholly contiguous nature of the signal in the Gershman et al. assembly, this arrangement is best understood as an intra-chromosomal rearrangement between *Manduca* and *Bombyx* leading to an incorrect synteny ordering. **Bottom:** Examining deviations from background levels of variation with pairwise difference in genetic diversity (π) between populations. Diversity is roughly equal between populations (near 0 difference) except within putatively inverted regions, in which both NC and KS have fewer pairwise differences than AZ. In both the top and bottom, scaffold limits are denoted by dashed vertical lines. **Left:** The majority of assembled scaffolds could be placed in an order with synteny. **Right:** Some scaffolds could not be placed by this method and appear here, unordered. Note that some among these scaffolds also show scaffold-wide differentiation and differences in diversity. Compare to Figure 4 of the main text, in which the full length of the putative inversion appears to have been contiguously assembled.

**Figure S3.**
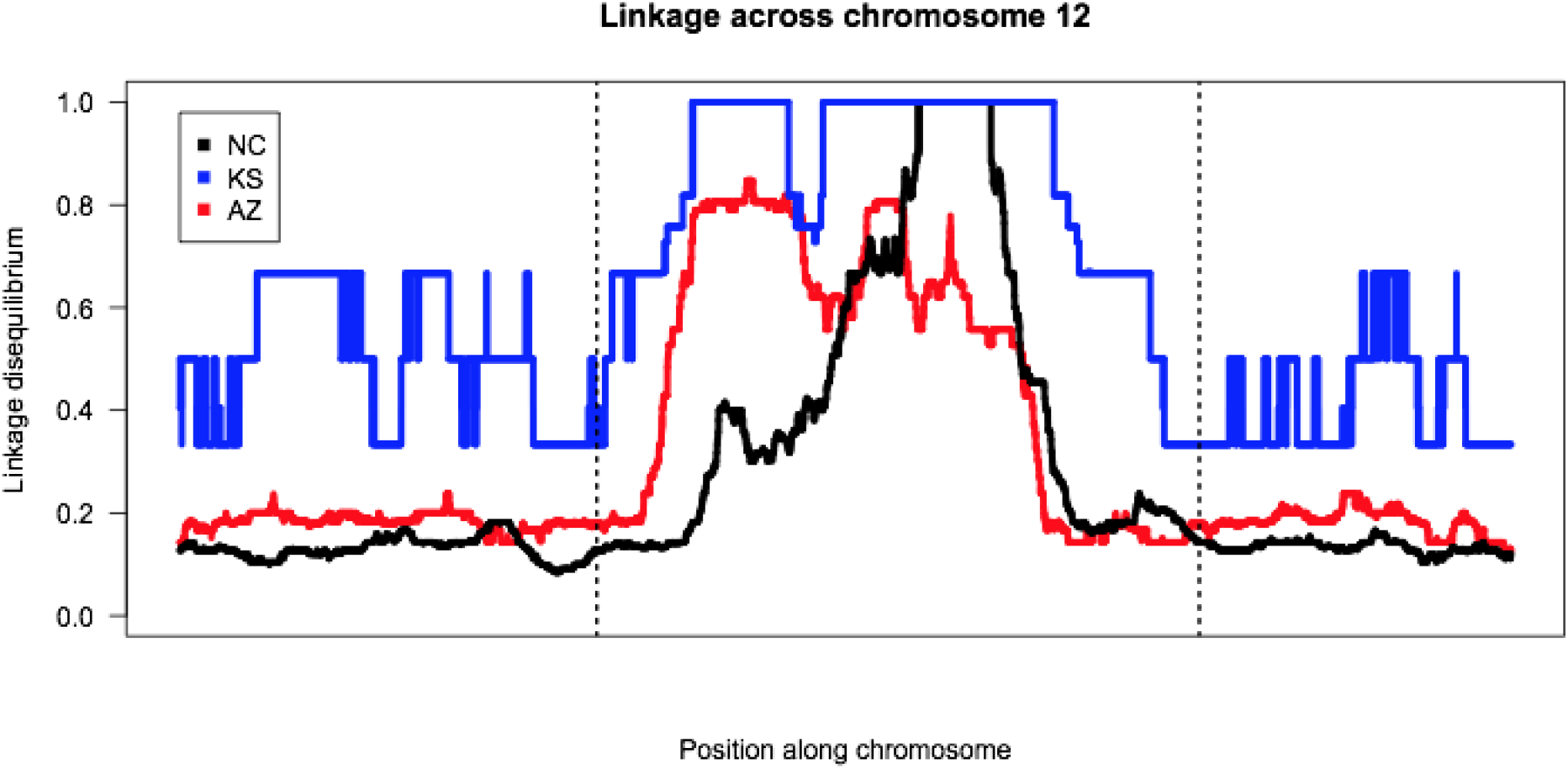
Linkage disequilibrium across chromosome 12. A rolling median of windows of 10Kb for the genotype ρ^2^ as calculated with vcftools in each population. Dashed lines indicate approximate location of the inferred inversion breakpoints. Note as well with the fewest sampled chromosomes in the Kansas population, apparent linkage is overall higher across the chromosome.

**Figure S4.**
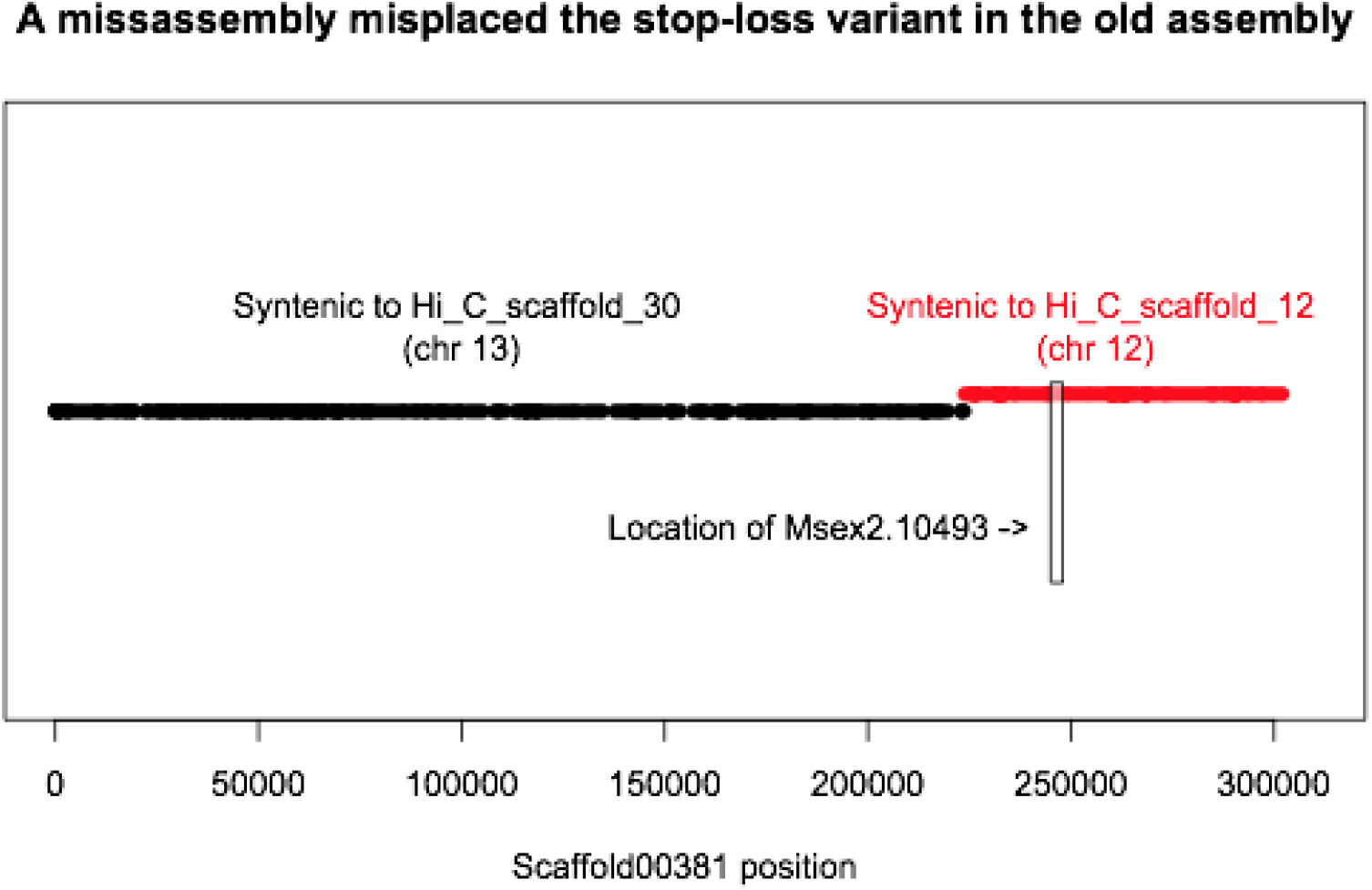
Syntenic alignment of scaffold00381 compared to the chromosome-level assembly. The majority of the scaffold (in black) is syntenic to chromosome 13, explaining its previous assignment to that chromosome. However, a significant portion of the sequence (red), including Msex2.10493, the putative segregating pseudogene, is syntenic to chromosome 12.

**Figure S5.**
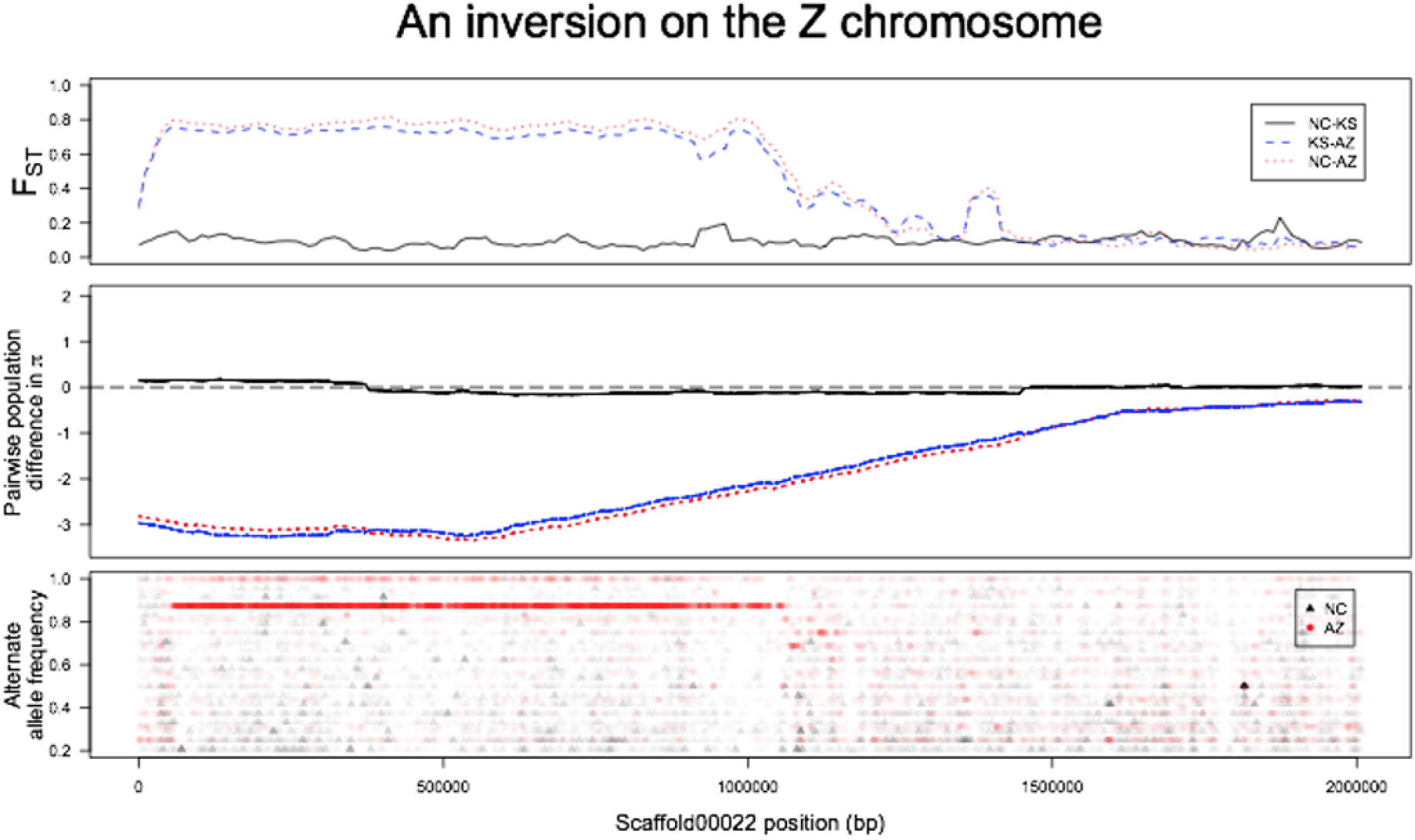
Evidence for an inversion on the Z chromosome using the Kanost et al. assembly. **Top.** A roughly 1 Mb portion of scaffold0002 (from 58 Kb to 1.03 Mb) shows elevated differentiation in North Carolina – Arizona and Kansas – Arizona comparisons. **Middle.** As with the chromosome 12 inversion, this region shows differences in π between the inverted and non-inverted region. **Bottom.** Like the chromosome 12 inversion, Arizona shows a long tract of shared allele frequencies in the region while North Carolina does not (note that Kansas allele frequencies are omitted due to lower sampling effort artificially stratifying frequencies).

**Figure S6.**
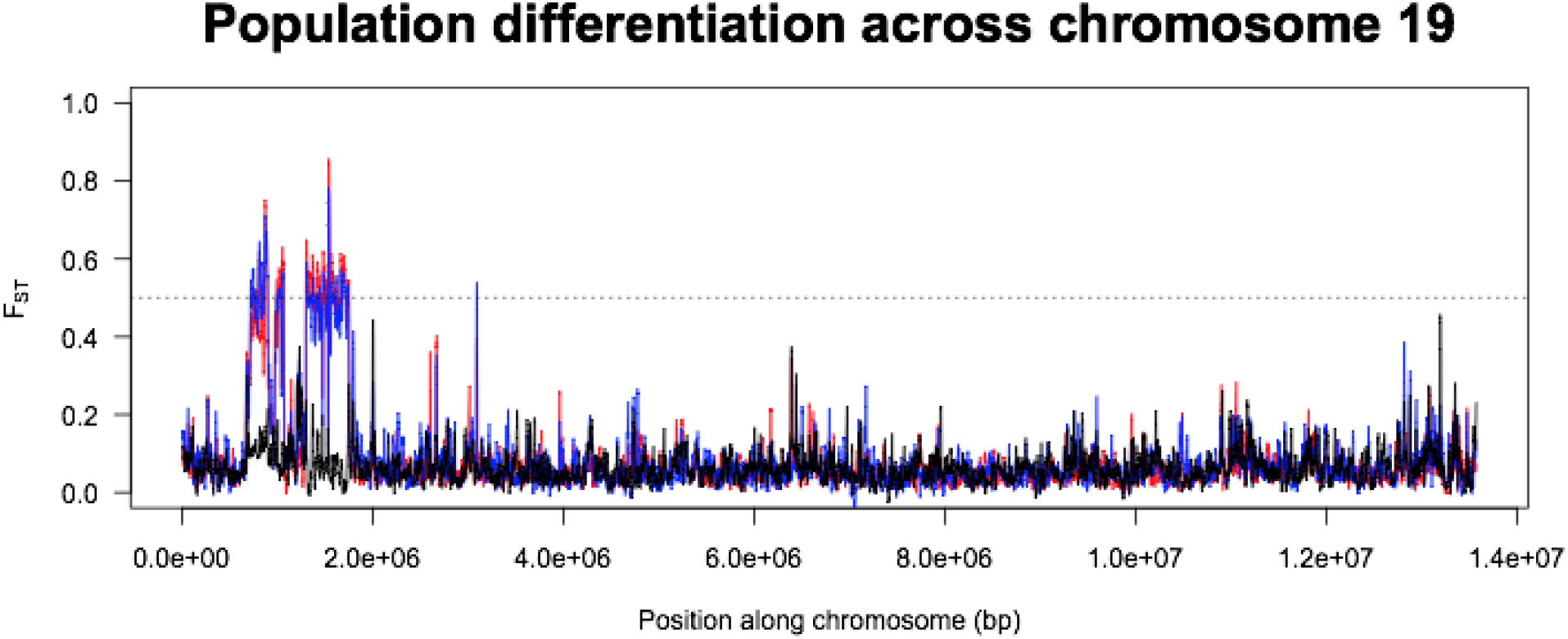
Population differentiation on chromosome 19. Two separate regions of elevated F_ST_ occur in the first two megabases of the chromosome, corresponding to the two different outlier scaffolds in the Kanost et al. assembly.

## References

Anderson, A.R., A.A. Hoffmann, S. W. McKechnie, P. A. Umina, and A.R. Weeks, 2005 The latitudinal cline in the In (3R) Payne inversion polymorphism has shifted in the last 20 years in Australian Drosophila melanogaster populations. Molecular Ecology 14: 851–858.

Banister, M. J., and R. H. White, 1987 Pigment migration in the compound eye of Manduca sexta: effects of light, nitrogen and carbon dioxide. Journal of insect physiology 33: 733–743.

Berry, J., P. Bretting, G. Nabhan, and C. Weber, 1981 Domesticated Proboscidea parviflora: a potential oilseed crop for arid lands. Journal of Arid Environments 4: 147–160.

Boer, G. de, and F. E. Hanson, 1987 Differentiation of roles of chemosensory organs in food discrimination among host and non-host plants by larvae of the tobacco hornworm, Manduca sexta. Physiological Entomology 12: 387–398.

Cao, X., and H. Jiang, 2017 An analysis of 67 RNA-seq datasets from various tissues at different stages of a model insect, *Manduca sexta*. BMC Genomics 18:.

Charlesworth, B., J. A. Coyne, and N. H. Barton, 1987 The relative rates of evolution of sex chromosomes and autosomes. The American Naturalist 130: 113–146.

Cingolani, P., A. Platts, L. L. Wang, M. Coon, T. Nguyen et al., 2012 A program for annotating and predicting the effects of single nucleotide polymorphisms, SnpEff: SNPs in the genome of *Drosophila melanogaster* strain w 1118; iso-2; iso-3. Fly 6: 80–92.

Connallon, T., C. Olito, L. Dutoit, H. Papoli, F. Ruzicka et al., 2018 Local adaptation and the evolution of inversions on sex chromosomes and autosomes. Philosophical Transactions of the Royal Society B: Biological Sciences 373: 20170423.

Contreras, H. L., J. Goyret, M. von Arx, C. T. Pierce, J. L. Bronstein et al., 2013 The effect of ambient humidity on the foraging behavior of the hawkmoth Manduca sexta. Journal of Comparative Physiology A 199: 1053–1063.

Danecek, P., A. Auton, G. Abecasis, C. A. Albers, E. Banks et al., 2011 The variant call format and VCFtools. Bioinformatics 27: 2156–2158.

Dobzhansky, T., and A. H. Sturtevant, 1938 Inversions in the chromosomes of Drosophila pseudoobscura. Genetics 23: 28.

Evanno, G., S. Regnaut, and J. Goudet, 2005 Detecting the number of clusters of individuals using the software STRUCTURE: a simulation study. Molecular ecology 14: 2611–2620.

Ferguson, D., 1991 An essay on the long-range dispersal and biogeography of Lepidoptera, with special reference to the Lepidoptera of Bermuda. Memoirs of the Entomological Society of Canada 158: 67–79.

Fuller, Z. L., S. A. Koury, N. Phadnis, and S. W. Schaeffer, 2019 How chromosomal rearrangements shape adaptation and speciation: Case studies in Drosophila pseudoobscura and its sibling species Drosophila persimilis. Molecular ecology 28: 1283–1301.

Gershman, A., T. G. Romer, Y. Fan, R. Razaghi, W. A. Smith et al., 2021 De novo genome assembly of the tobacco hornworm moth (Manduca sexta). G3 11: jkaa047.

Grabherr, M. G., P. Russell, M. Meyer, E. Mauceli, J. Alföldi et al., 2010 Genome-wide synteny through highly sensitive sequence alignment: Satsuma. Bioinformatics 26: 1145–1151.

Haber, W. A., and G. W. Frankie, 1989 A tropical hawkmoth community: Costa Rican dry forest Sphingidae. Biotropica 155–172.

Höglund, G., and G. Struwe, 1970 Pigment migration and spectral sensitivity in the compound eye of moths. Zeitschrift für vergleichende Physiologie 67: 229–237.

Janzen, D. H., 1984 Two ways to be a tropical big moth: Santa Rosa saturniids and sphingids.

Kanost, M. R., E. L. Arrese, X. Cao, Y.-R. Chen, S. Chellapilla et al., 2016 Multifaceted biological insights from a draft genome sequence of the tobacco hornworm moth, *Manduca sexta*. Insect Biochemistry and Molecular Biology 76: 118–147.

Kanost, M. R., H. Jiang, and X. Q. Yu, 2004 Innate immune responses of a lepidopteran insect, *Manduca sexta*. Immunological Reviews 198: 97–105.

Kapun, M., D. K. Fabian, J. Goudet, and T. Flatt, 2016 Genomic evidence for adaptive inversion clines in Drosophila melanogaster. Molecular biology and evolution 33: 1317–1336.

Kawahara, A. Y., J. W. Breinholt, F. V. Ponce, J. Haxaire, L. Xiao et al., 2013 Evolution of Manduca sexta hornworms and relatives: Biogeographical analysis reveals an ancestral diversification in Central America. Molecular Phylogenetics and Evolution 68: 381–386.

Kirkpatrick, M., and N. Barton, 2006 Chromosome inversions, local adaptation and speciation. Genetics 173: 419–434.

Knief, U., G. Hemmrich-Stanisak, M. Wittig, A. Franke, S. C. Griffith et al., 2016 Fitness consequences of polymorphic inversions in the zebra finch genome. Genome biology 17: 199.

Kopelman, N. M., J. Mayzel, M. Jakobsson, N. A. Rosenberg, and I. Mayrose, 2015 Clumpak: a program for identifying clustering modes and packaging population structure inferences across K. Molecular ecology resources 15: 1179–1191.

Korneliussen, T. S., A. Albrechtsen, and R. Nielsen, 2014 ANGSD: Analysis of Next Generation Sequencing Data. BMC Bioinformatics 15:.

Kryuchkova-Mostacci, N., and M. Robinson-Rechavi, 2017 A benchmark of gene expression tissue-specificity metrics. Briefings in Bioinformatics 18: 205–214.

Lande, R., 1984 The expected fixation rate of chromosomal inversions. Evolution 743–752.

Langmead, B., and S. L. Salzberg, 2012 Fast gapped-read alignment with Bowtie 2. Nat Methods 9: 357–359.

Lasne, C., C. M. Sgrò, and T. Connallon, 2017 The Relative Contributions of the X Chromosome and Autosomes to Local Adaptation. Genetics 205: 1285–1304.

Lee, Y. W., L. Fishman, J. K. Kelly, and J. H. Willis, 2016 A segregating inversion generates fitness variation in yellow monkeyflower (Mimulus guttatus). Genetics 202: 1473–1484.

Lou, R. N., A. Jacobs, A. Wilder, and N. O. Therkildsen, 2020 A beginner’s guide to low-coverage whole genome sequencing for population genomics.

Lunter, G., and M. Goodson, 2011 Stampy: A statistical algorithm for sensitive and fast mapping of Illumina sequence reads. Genome Research 21: 936–939.

Lyons, J. I., A. A. Pierce, S. M. Barribeau, E. D. Sternberg, A. J. Mongue et al., 2012 Lack of genetic differentiation between monarch butterflies with divergent migration destinations. Molecular Ecology 21: 3433–3444.

McKenna, A. H., M. Hanna, E. Banks, A. Sivachenko, K. Cibulskis et al., 2010 The Genome Analysis Toolkit: A MapReduce framework for analyzing next-generation DNA sequencing data. Genome Research 20: 1297–1303.

McNeil, J., 1995 Physiological integration of migration in Lepidoptera. Insect migration: tracking resources through space and time.

Mechaber, W. L., and J. G. Hildebrand, 2000 Novel, non-solanaceous hostplant record for Manduca sexta (Lepidoptera: Sphingidae) in the Southwestern United States. Annals of the Entomological Society of America 93: 447–451.

Meisner, J., and A. Albrechtsen, 2018 Inferring population structure and admixture proportions in low-depth NGS data. Genetics 210: 719–731.

Mérot, C., E. Berdan, H. Cayuela, H. Djambazian, A.-L. Ferchaud et al., 2021 Locally adaptive inversions modulate genetic variation at different geographic scales in a seaweed fly. bioRxiv 2020–12.

Mongue, A. J., M. E. Hansen, L. Gu, C. E. Sorenson, and J. R. Walters, 2019 Nonfertilizing sperm in Lepidoptera show little evidence for recurrent positive selection. Molecular Ecology 28: 2517–2530.

Mongue, A. J., M. E. Hansen, and J. R. Walters, 2021 Support for faster and more adaptive Z chromosome evolution in two divergent lepidopteran lineages. Evolution n/a:

Mongue, A. J., M. E. Hansen, and J. R. Walters, 2020 ZZ Top: faster and more adaptive Z chromosome evolution in two Lepidoptera. bioRxiv.

Mongue, A. J., and J. Walters, 2017 The Z chromosome is enriched for sperm proteins in two divergent species of Lepidoptera. Genome 61: 248–253.

Nardi, J. B., 1993 Modulated expression of a surface epitope on migrating germ cells of Manduca sexta embryos. Development 118: 967–975.

Nei, M., 1987 Molecular evolutionary genetics. Columbia university press.

Nei, M., Y. Niimura, and M. Nozawa, 2008 The evolution of animal chemosensory receptor gene repertoires: roles of chance and necessity. Nature Reviews Genetics 9: 951–963.

Prieto-Godino, L. L., R. Rytz, B. Bargeton, L. Abuin, J. R. Arguello et al., 2016 Olfactory receptor pseudo-pseudogenes. Nature 539: 93–97.

R Core Team, 2017 R Core Team (2017). R: A language and environment for statistical computing. R Foundation for Statistical Computing, Vienna, Austria. URL http://www.R-project.org/. R Foundation for Statistical Computing.

Rausch, T., T. Zichner, A. Schlattl, A. M. Stütz, V. Benes et al., 2012 DELLY: structural variant discovery by integrated paired-end and split-read analysis. Bioinformatics 28: i333–i339.

Rice, W. R., 1984 Sex chromosomes and the evolution of sexual dimorphism. Evolution 38: 735–742.

Shapiro, A. M., 2006 Survival of freezing and subsequent summer eclosion by three migratory moths: Manduca sexta and Hyles Lineata (Sphingidae), and Helicoverpa zea (Noctuidae). Journal of the Lepidopterists’ Society 60: 101–103.

Skotte, L., T. S. Korneliussen, and A. Albrechtsen, 2013 Estimating individual admixture proportions from next generation sequencing data. Genetics 195: 693–702.

Stevenson, R., K. Corbo, L. Baca, and Q. Le, 1995 Cage size and flight speed of the tobacco hawkmoth Manduca sexta. Journal of Experimental Biology 198: 1665–1672.

Sturtevant, A. H., 1917 Genetic factors affecting the strength of linkage in Drosophila. Proceedings of the National Academy of Sciences of the United States of America 3: 555.

The International Silkworm Genome Consortium, 2008 The genome of a lepidopteran model insect, the silkworm *Bombyx mori*. Insect Biochemistry and Molecular Biology 38: 1036–1045.

Todesco, M., G. L. Owens, N. Bercovich, J.-S. Légaré, S. Soudi et al., 2020 Massive haplotypes underlie ecotypic differentiation in sunflowers. Nature 584: 602–607.

Wagner, D. L., 2005 Caterpillars of eastern North America: a guide to identification and natural history. Princeton University Press.

Wellenreuther, M., H. Rosenquist, P. Jaksons, and K. W. Larson, 2017 Local adaptation along an environmental cline in a species with an inversion polymorphism. Journal of Evolutionary Biology 30: 1068–1077.

Whittington, E., Q. Zhao, K. Borziak, J. R. Walters, and S. Dorus, 2015 Characterisation of the *Manduca sexta* sperm proteome: Genetic novelty underlying sperm composition in Lepidoptera. Insect Biochemistry and Molecular Biology 62: 183–193.

Wysoker, A., K. Tibbetts, and T. Fennell, 2013 Picard tools version 1.90. http://picard. sourceforge. net (Accessed 14 December 2016) 107: 308.

Yasukochi, Y., M. Tanaka-Okuyama, F. Shibata, A. Yoshido, F. Marec et al., 2009 Extensive conserved synteny of genes between the karyotypes of *Manduca sexta* and *Bombyx mori* revealed by BAC-FISH mapping. PLoS ONE 4: e7465:

Zhan, S., W. Zhang, K. Niitepõld, J. Hsu, J. F. Haeger et al., 2014 The genetics of monarch butterfly migration and warning colouration. Nature 514: 317–321.

Zhang, Z.-J., S.-S. Zhang, B.-L. Niu, D.-F. Ji, X.-J. Liu et al., 2019 A determining factor for insect feeding preference in the silkworm, Bombyx mori. PLOS Biology 17: e3000162.

